# Recombinant Fsh and Lh therapy for spawning induction of previtellogenic and early spermatogenic arrested teleost, the flathead grey mullet (*Mugil cephalus*)

**DOI:** 10.1101/2021.09.29.462352

**Authors:** Sandra Ramos-Júdez, Ignacio Giménez, Josep Gumbau-Pous, Lucas Stephen Arnold-Cruañes, Alicia Estévez, Neil Duncan

## Abstract

With the expansion and diversification of global aquaculture, efforts continue to develop new bio-technologies for assisted reproduction in species that present reproductive dysfunctions. Flathead grey mullet (*Mugil cephalus*) held in intensive conditions in the Mediterranean region, display a severe reproductive dysfunction, where males do not produce fluent milt and females are arrested at previtellogenesis or early stages of vitellogenesis. In the present study, weekly injections of species-specific single-chain recombinant gonadotropins (rGths); follicle stimulating hormone (rFsh) (6 to 12 μg kg^-1^ doses) and luteinizing hormone (rLh) (2.5 to 24 μg kg^-1^ doses) were administered to induce vitellogenesis, from previtellogenesis / early vitellogenesis to the completion of vitellogenic growth in females and enhance spermatogenesis to produce adequate volumes of sperm from non-fluent males. During the experiment, all treated females (n = 21) developed oocytes in late vitellogenesis with 603 ± 8 μm diameter and all treated males produced fluent sperm. To induce oocyte maturation, ovulation and spawning, females were treated with either (i) a priming dose of 30 μg kg^-1^ of rLh and a resolving dose of 40 mg kg^-1^ of progesterone (P_4_), (ii) priming and resolving doses of 30 μg kg^-1^ of rLh, or (iii) priming and resolving doses of 40 mg kg^-1^ of P_4_ given 24:05 ± 0:40 h apart. Females were placed in spawning tanks with rGth treated males that had fluent sperm. Spontaneous spawns of fertilised eggs were obtained after inducing with rLh + P_4_ or rLh + rLh (priming and resolving injections) with a spawning success of the 85% (8 of 9 females) and 100% (n = 6), respectively. The eggs collected from the tanks presented 64 ± 22% fertilization with embryo development and 57 ± 24 % hatching. The treatment P_4_ + P_4_ had a lower ovulation success (50 % - 3 of 6 females) and spawning success (17 %) with no fertilised eggs. Success was independent of the initial gonadal stage of females. In comparison, control females did not show any advance in gonadal development from initial stages and control males did not produce fluent sperm. The present results confirm the possibility of controlling oogenesis from previtellogenesis to the completion of maturation and fertilised tank spawning using exclusively rFsh and rLh in a teleost species.

## 1. Introduction

Intensive aquaculture is looking for ways to improve reproductive control, especially in reproductively dysfunctional species to ensure the supply of fry for large-scale commercial production. The development of culture protocols will not only ensure a consistent and sustainable supply for grow-out operations, but will also allow for genetic improvements through selective breeding. The success of culture protocols, will in turn alleviate the fishery pressure on stocks of natural populations that in many cases are compromised. An essential part to provide a reliable supply of juveniles is to control the reproduction of adult fish held in captivity. However, some species do not complete reproduction in captivity and exogenous hormonal therapies have been employed to develop aquaculture production.

The two reproductive stages, gametogenesis (oogenesis and spermatogenesis) and maturation (oocyte maturation and spermiation) are controlled by different reproductive hormones produced in the pituitary and gonad, i.e., gonadotropin hormones (Gths) and steroids (Mañanós et al., 2009). Hormone therapies based on gonadotropin releasing hormones and luteinizing hormone receptor agonists (human chorionic gonadotropins or pituitary extracts) are commonly used to control the maturation phase, while hormonal control of gametogenesis is rarely used in the aquaculture industry (Mylonas and Zohar, 2007). The use of relatively new recombinant gonadotropin hormones (rGths), the recombinant follicle-stimulating (rFsh) and luteinizing hormones (rLh), can open new strategies in aquaculture to treat reproductive disorders and develop out-of-season breeding programs (Molés et al., 2020). To this end, different *in vivo* treatments have been developed, mainly focused on final maturation and spermiation/ovulation stages by single or double rGths injections (Aizen et al., 2017; Kobayashi et al., 2006). However, fish species arrested at the early stages of the reproductive cycle require control of gametogenesis, with long-term treatments of repeated injections that maintain elevated plasma levels of specific Gths (Molés et al., 2020). In the case of males, different successful long-term approaches have been described for immature European eel (*Anguilla anguilla*) (Peñaranda et al., 2018) and mature Senegalese sole (*Solea senegalensis*) (Chauvigné et al., 2018, 2017). In the case of females, it has been difficult to define similar long-term treatments to produce viable gametes from females arrested prior to vitellogenesis. A significant advance was achieved with the long-term treatment of previtellogenic flathead grey mullet (*Mugil cephalus*) females with rFsh and rLh to successfully complete vitellogenesis (Ramos-Júdez et al., 2021). However, after maturation induction with rLh and Progesterone (P_4_), females held with spermiating males failed to spawn spontaneously. Therefore, gametes were stripped and artificially fertilised and a low percentage of fertilisation (<1%) was obtained, which questioned the viability of the process for aquaculture purposes. However, despite of the low fertilisation, the study demonstrated the possibility of using rFsh and rLh to induce oogenesis from previtellogenesis to obtain eggs and larvae in intensive conditions and encouraged further research to improve the results obtained.

The flathead grey mullet, has a worldwide distribution in tropical, subtropical and temperate waters (McDonough et al., 2005), tolerance to wide ranges of salinities, excellent flesh quality and high growth rates. It represents an important species and a potential candidate in the diversification of aquaculture products mainly in the Mediterranean area, the Southeast of Asia, Taiwan, Japan and Hawaii (González-Castro and Minos, 2016). In the Mediterranean, flathead grey mullet spawn from July to October (Whitfield et al., 2012), however, breeders held in intensive conditions show reproductive dysfunctions; males rarely produce fluent milt (Aizen et al., 2005; De Monbrison et al., 1997; Yashouv, 1969) and females are found to be arrested at previtellogenesis (Ramos-Júdez et al., 2021) or early stages of vitellogenesis (Aizen et al., 2005).

The objective of the present study was to demonstrate that long-term treatment with rGths, rFsh and rLh, can induce gametogenesis in flathead grey mullet males arrested with no fluent spermiation and females arrested at early stages of gametogenesis to obtain spawning of eggs with good fertilisation rates that provide viable larvae. Different treatments based on rLh and P_4_ were tested to induce oocyte maturation and spawning in females. As a final step, to check larval development and growth, the eggs obtained were used to carry out a preliminary larval rearing trial using a mesocosm technique.

## 2. Material and methods

### 2.1 Broodstock maintenance

A flathead grey mullet broodstock was formed with individuals, originally obtained from the Ebro River (Spain) or from a semi-extensive pond fish farm (Finca Veta La Palma, Isla Mayor, Spain) and which had been held for 1.5 to 3.5 years in IRTA facilities (Sant Carles de la Ràpita, Spain). Thirty females with weights ranging from 0.9 to 2.4 kg and standard length (SL) from 37 to 53 cm, and fifteen males ranging from 0.7 to 1.3 kg and 34 to 43.5 cm were used. All fish were larger than the reported SL for first maturation in this species (27 - 35 cm for females, 25 - 30 cm for males) (Whitfield et al., 2012). To identify individuals each fish was tagged intramuscularly with a Passive Integrated Transponder (PIT) tag (Trovan®, ZEUS Euroinversiones S.L. Madrid, Spain). The sex of the individuals was determined by the presence or absence of oocytes obtained through slight suction with a 1.67-mm plastic catheter inserted through the genital opening. Fish were maintained in 10 m^3^ covered tanks in a recirculating system (IRTAmar®) supplied with 36 ‰ salinity water under natural conditions of light and controlled winter temperatures (≥ 14°C) during the last year. Before the study, conducted from early August to early November, the selected broodstock was transferred to another 10 m^3^ tank, water temperature was controlled at 23.1 ± 0.2 °C and photoperiod was ambient (14L:10D - 11L:13D). Fish were fed 5 days a week at the rate of 1.5% of their body weight with a mix of two commercial marine fish diets; 90% mix of Le-2 and Le-5 Europa RG (Skretting, Spain) and 10% Brood Feed Lean (Sparos, Portugal). During all experimental procedures, for hormone administration and sampling, fish were anaesthetised with 73 mg L^-1^ of MS-222. When required for the study, males were euthanised with an overdose of MS-222 (250 mg L^-1^) and death was confirmed by a cut in the gills to exsanguinate the fish.

### 2.2 Hormonal induction

#### 2.2.1. Induction of vitellogenesis

Females were assigned randomly to rGth and control groups taking care to have similar distribution of females in different initial maturation stages in both groups. The control group (total n = 9) was formed with 6 females in previtellogenesis (5 in primary growth and 1 in cortical alveoli step) and 3 in early vitellogenesis. A total of 21 females were assigned to receive the hormonal treatment; 12 females initially were in previtellogenesis (8 in primary growth and 4 in cortical alveoli stage) and 9 in early vitellogenesis. Those females in early vitellogenesis had longer time in intensive captive conditions (> 2.25 years) although not all females that were held for this time had started vitellogenesis.

Single chain recombinant *Mugil cephalus* rFsh and rLh produced in Chinese hamster ovary (CHO) cells were purchased from Rara Avis Biotec S.L. (Valencia, Spain). *Mugil cephalus* rFsh was supplied with a concentration of 12 μg mL^-1^ and rLh with concentration of 8 μg mL^-1^. A methodology based on the protocol described by Ramos-Júdez *et al*. (2021) was followed. The pattern of application of rFsh and rLh aimed to mimic the physiological variations of Fsh and Lh associated with natural reproductive development; initially only rFsh was administered during the early stages of gametogenesis and, subsequently, a decrease in rFsh with an increase of rLh to regulate late gametogenesis (Lubzens et al., 2010). The protocol was applied according to ovarian development (Fig 1). Increasing weekly doses of 6, 9 and 12 μg kg^-1^ rFsh were administered intramuscularly to induce previtellogenesis (~100 μm oocyte diameter) to vitellogenesis (> 200 μm). Weekly doses were maintained at 12 μg kg^-1^ during vitellogenesis. As vitellogenesis progressed and when mean diameter of the most developed oocytes was ≥ 300 μm, females received in addition a weekly administration of rLh at rising doses. A dose of 2.5 μg kg^-1^ was maintained until females entered into late-vitellogenesis (≥ 400 μm) (Greeley et al., 1987) and was then increased to 4 and 6 μg kg^-1^. At an oocyte diameter of ~ 500 μm, weekly rFsh doses were reduced to 4 μg kg^-1^ while rLh was increased to 9 μg kg^-1^. A combination of 4 μg kg^-1^ rFsh and 12 μg kg^-1^ rLh per week was administered until vitellogenic growth was completed. The completion of oocyte growth was determined when oocytes were deemed approaching maturation; microscopic examination showed that the most developed oocytes were nearing 600 μm in diameter. The nine females in the control group were also manipulated each week and were injected with saline solution (1 mL) a total of twelve times.

**Figure 1.**
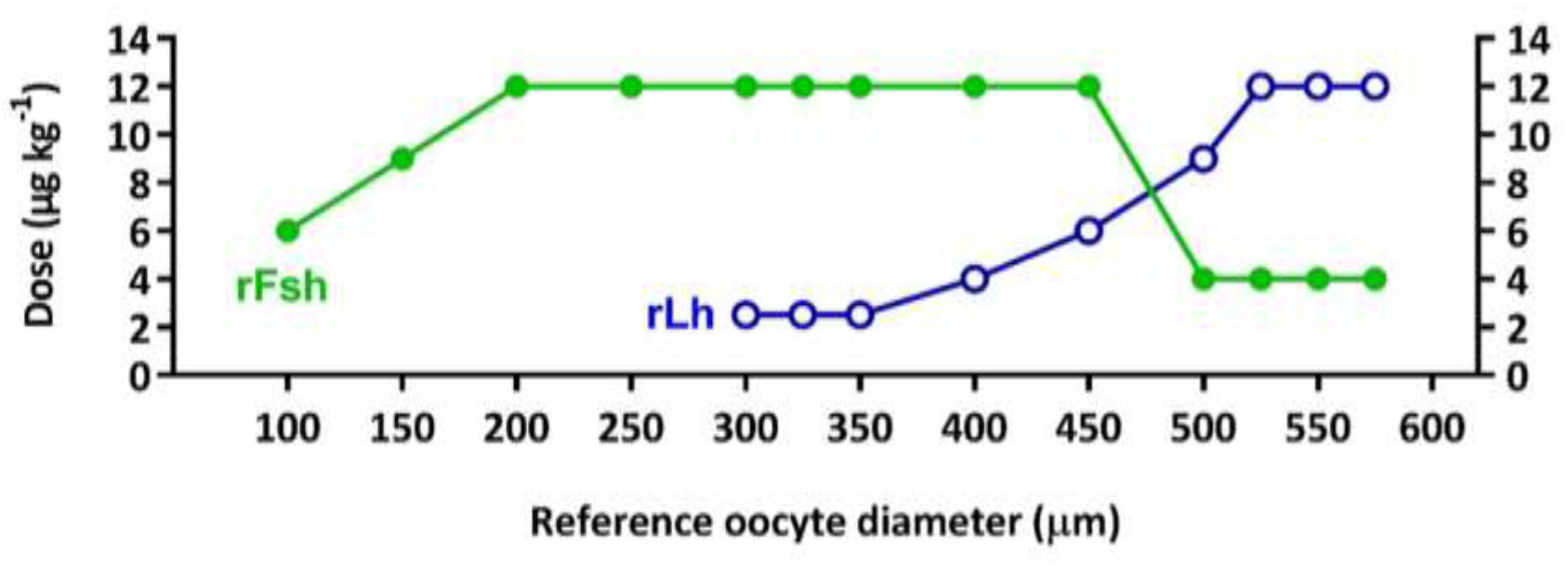
Schematic representation of the rFsh and rLh treatment applied to flathead grey mullet females (n = 21). The protocol was applied according to the development of female gonads determined by ovarian biopsies. Weekly doses of 6, 9 and 12 μg kg-1 rFsh were applied to induce previtellogenesis (~100 μm oocyte diameter) to vitellogenesis (> 200 μm) and vitellogenic growth was maintained with a dose of 12 μg kg-1 per week. When the mean diameter of the largest oocytes was ≥ 300 μm, in addition to rFsh, rLh was administered in increasing doses. A dose of 2.5 μg kg-1 was maintained until females presented ≥ 400 μm oocytes and was raised to 4 and 6 μg kg-1 rLh. At ~500 μm diameter, weekly rFsh doses were reduced to 4 μg kg-1 whereas rLh was increased to 9 μg kg-1. A combination of 4 μg kg-1 rFsh and 12 μg kg-1 rLh per week were administered until vitellogenic growth was completed (~600 μm). Each point corresponds to a weekly administration. This scheme represents the longest pattern of administration of those females that required a total of thirteen weeks to complete vitellogenic growth.

Assessment of gonadal development was undertaken on alternate weeks by ovarian biopsies obtained by slight suction through a plastic cannula. Fresh ovarian samples were examined under a microscope (×40 magnification), to measure the mean diameter of the largest most advanced oocytes (n = 20 per female) and a sample fixed for histology. Blood samples were obtained before the initial treatment (week 0) and when vitellogenic growth was completed in the treated group and at the end of the experiment in the control group.

#### 2.2.2. Hormonal administration in males

In parallel, males were assigned randomly to rGth and control groups taking care to have similar distribution of males in different initial maturation stages in both groups. The control group (total n = 6) was formed with 5 males with no presence of sperm and 1 with a presence of sperm (sperm index 1, presence of sperm, but not fluid, see below). A total of 9 males were assigned to receive the hormonal treatment; 7 males with no presence of sperm and 2 with presence of sperm. Males treated with rGth were split into two groups that received the same treatment, but at different times in the experimental period. The reason was to assure the availability of spermiating males for spawning induction when the females completed vitellogenic growth. Group 1 of males (n = 4 treated, n = 3 control) initiated the treatment on week 1 and the group 2 (n = 5 treated, n = 3 control) on week 3 (Fig 2). The aim of the treatment was to apply rGths according to their described role in spermatogenesis. Increasing doses of rFsh (6, 9 and 12 μg kg^-1^) were administered at early spermatogenesis and high levels (12 μg kg^-1^) during testicular growth (Fig 3). No rLh was administered during early spermatogenesis, both rFsh and rLh (12 μg kg^-1^ of both rFsh and rLh) during the middle stages of testicular growth and only rLh (12 μg kg^-1^) was administered at late stages to induce sperm maturation (Schulz et al., 2010). Group 1 received the treatment for a total of 12 weeks whereas group 2 for 10 weeks. At week 9, group 1 received a dose of 12 μg kg^-1^ rFsh instead of rLh as a reduction in the spermiation stage was observed (see 3.2 section). Doses were in the range of previous studies with males using rGths produced in CHO cells (Chauvigné et al., 2018, 2017; Peñaranda et al., 2018; Ramos-Júdez et al., 2021). Control males were injected with 1 mL of saline solution per week.

**Figure 2.**
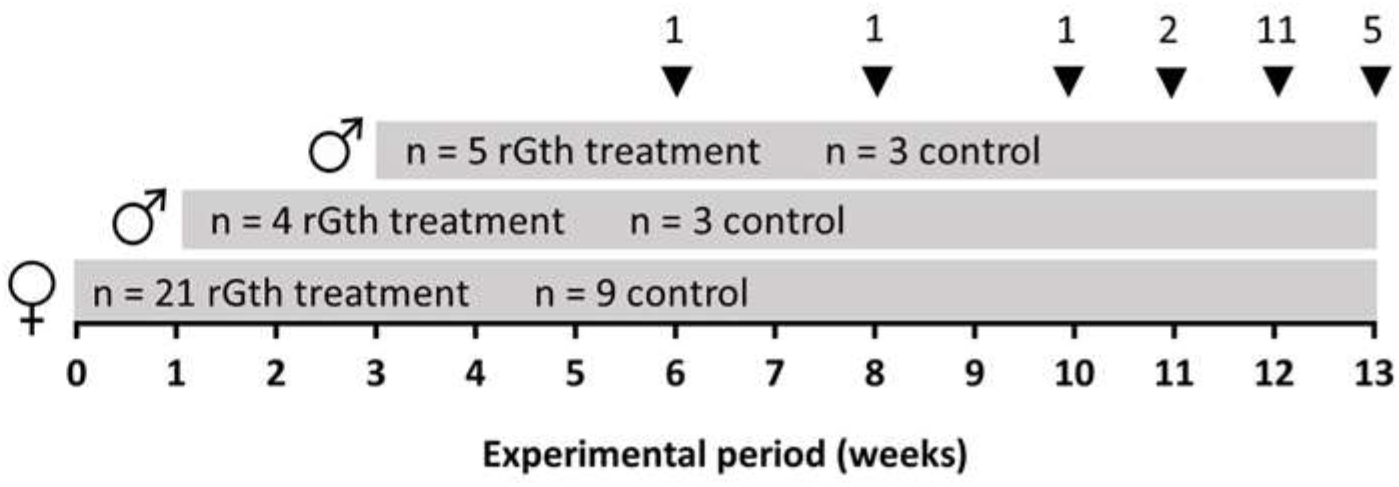
Overall scheme of the experiment. Females receiving the rGths treatment were injected weekly until completion of vitellogenic growth which took place in a maximum period of thirteen weeks, following thirteen administrations. Arrow heads indicate the moment in which individual females were induced to spawn (number of females indicated by the number on the top). Control females received a total of twelve saline injections. Group 1 of males started the treatments (rGth treatment and saline) on week 1, whereas the group 2 on week 3.

**Figure 3.**
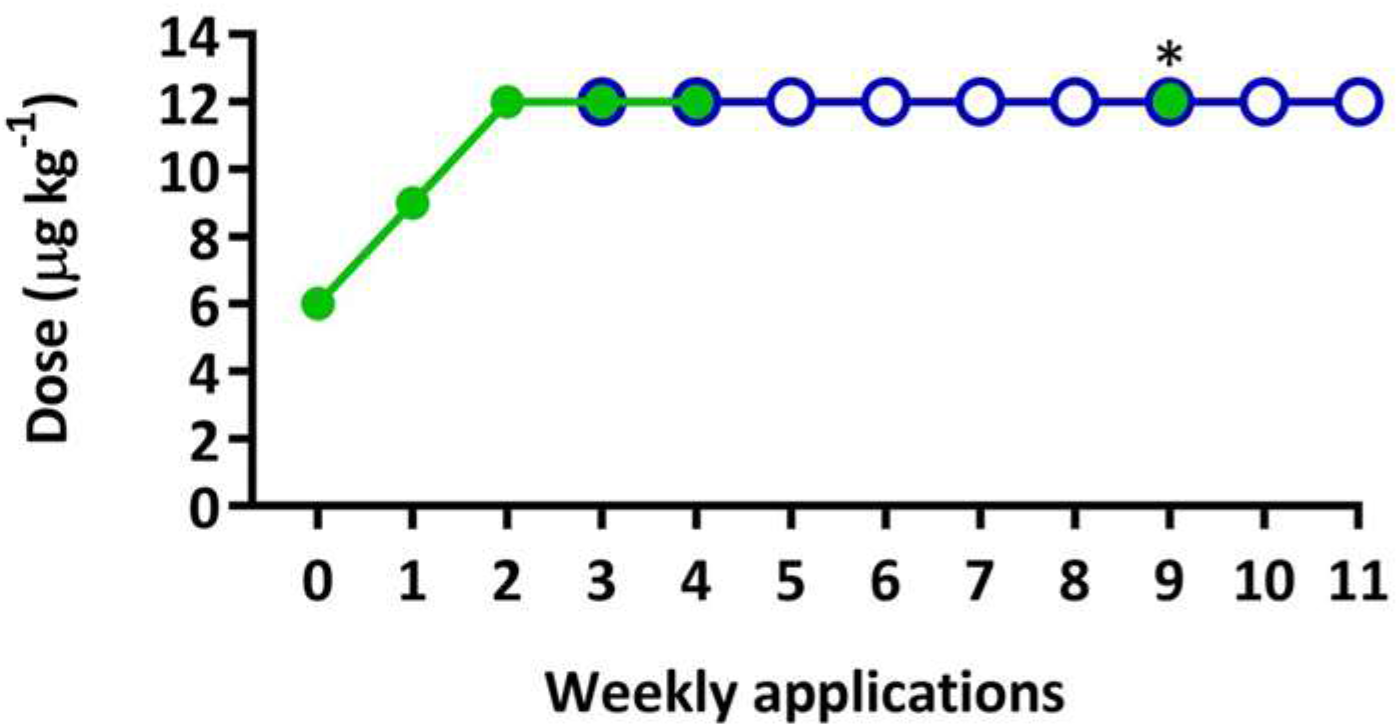
Diagram of the hormonal treatment of flathead grey mullet males. Asterisk indicates the moment that group 1 received 12 μg kg-1 of rFsh instead of rLh. Green indicates rFsh administration and blue indicates rLh administration.

In order to evaluate the progression of maturation, at the beginning of the treatment and on a weekly basis, a gentle squeeze on the ventral abdomen toward the urogenital opening was applied to release the sperm. Spermiation stage was determined on a scale from 0 to 3 (0 = not fluent, 1 =fluent but no sample can be obtained, 2 = fluent, 3 = very fluent). By this method, only mature cells are released together with the seminal plasma. Therefore, to determine the stage of development in males, some males were sacrificed at the beginning (n = 2) and the end of the treatment (n = 2 per group) and the testes removed for histological procedures and the measurement of gonadosomatic index (GSI: testes weight/fish weight x 100). Towards the end of the experiment (weeks 10 and 13), sperm samples from running males were collected, total volume recorded and stored at 4°C for sperm quality analysis. Blood samples were also taken from treated and control males each two weeks of the treatment.

#### 2.2.3. Oocyte final maturation and spawning induction

Three different treatments were followed for maturation and spawning induction. Females that completed vitellogenic growth (as determined by gonadal biopsy) and males with running milt (spermiating stage 2 or 3) were selected for spawning induction. The individuals were separated from the main group and stocked in a separate 10-m^3^ tank per treatment. The three separate tanks had the same conditions as the holding tank. One male in each spawning tank received a dose of 24 μg kg^-1^ of rLh while the others followed the hormonal treatment previously described, receiving 12 μg kg^-1^ of rLh. The selected females were injected intramuscularly with either (i) priming rLh and resolving Progesterone (P_4_) (Prolutex, IBSA Group, Italy) as in Ramos-Júdez *et al*. (2021), (ii) rLh for priming and resolving injections or (iii) P_4_ for priming and resolving injections (doses in Table 1). Priming and resolving injections were administered 24:05 ± 0:40 h apart. Ovarian samples were taken with a cannula and examined before each administration. Maturation and ovulation induction of females from each treatment group was staggered on different consecutive days in order to separate the different spawning events.

**Table 1.** Treatments to induce maturation and spawning in flathead grey mullet (*Mugil cephalus*) females and the results obtained. Latency period indicates the time (h) between resolving injection and spawning; Ovulation success (%) indicates the number of females that ovulated after the resolving injection divided by the total number of injected females; Spawning success (%) indicates the number of females that spawned naturally in the tank after the resolving injection, divided by the total number of injected females. Data are expressed as mean ± standard deviation. Oocyte diameter, total eggs per female and fecundity between the three treatments were compared by one-way ANOVA while latency period, percentages of initial fertilization, embryo survival, hatching and hatching in 96-well plate between the successful treatments (rLh + rLh and rLh + P_4_) were compared through a t-student test.

| Group | rLh + P<sub>4</sub> | rLh + rLh | P<sub>4</sub> + P<sub>4</sub> |
| --- | --- | --- | --- |
| Treatment Priming injection | 30 μg kg<sup>-1</sup> rLh | 30 μg kg<sup>-1</sup> rLh | 40 mg kg<sup>-1</sup> P<sub>4</sub> |
| Treatment Resolving injection | 40 mg kg<sup>-1</sup> P<sub>4</sub> | 30 μg kg<sup>-1</sup> rLh | 40 mg kg<sup>-1</sup> P<sub>4</sub> |
| Oocyte diameter (μm) | 606 ± 10<sup>a</sup> | 602 ± 6<sup>a</sup> | 600 ± 7<sup>a</sup> |
| No. of injected females | 9 | 6 | 6 |
| No. of spawned females | 8 | 6 | 1 |
| Latency period (h) | 16:36 ± 1:56<sup>a</sup> | 16:51 ± 2:22<sup>a</sup> | 24:30:00 |
| Ovulation success (%) | 100 | 100 | 50 |
| Spawning success (%) | 89 | 100 | 17 |
| Total eggs per female | 1,760,100 ± 821,102<sup>a</sup> | 1,631,889 ± 954,138<sup>a</sup> | 1,280,833 ± 966,42<sup>a</sup> |
| Fecundity (egg kg<sup>-1</sup> bw) | 1,194,637 ± 635,747<sup>a</sup> | 1,205,1387 ± 487,002<sup>a</sup> | 694,529 ± 526,457<sup>a</sup> |
| Initial fertilization (%) | 49 ± 23<sup>a</sup><br>(8 min – 80 max) | 60 ± 16<sup>a</sup><br>(39 min – 84 max) | 0 |
| Embryo survival (%) | 56 ± 22<sup>a</sup><br>(15 min – 80 max) | 73 ± 21<sup>a</sup><br>(42 min – 90 max) | 0 |
| Hatching (%) | 57 ± 21<sup>a</sup><br>(25 min – 88 max) | 61 ± 30<sup>a</sup><br>(23 min – 90 max) | - |
| Hatching 96-well plate (%) | 95 ± 4<sup>a</sup> | 97 ± 1<sup>a</sup> | - |

The sex ratio was 1:2 or 1:3 (female : male) per spawning event depending on availability of males. After all the spawning events, males were returned back with the main group to the initial tank. In the cases where females ovulated and showed a swollen belly, but did not release the eggs, manual stripping was applied.

Parallelly, oocytes obtained from cannulation from females (n = 6) that had received the priming injection of rLh (30 μg kg^-1^) were incubated *in vitro* with different hormones. A total of 58.7 ± 29.9 oocytes were incubated in a well of a 96-well plate with 200 μL of Leibovitz’s L-15 medium, with either no hormone (control), rLh (concentrations of 400, 200, 100, 50 and 10 ng mL^-1^), rFsh (400, 200, 100, 50 and 10 ng mL^-1^) or P_4_ (4000, 1000, 500 and 50 ng mL^-1^). Each treatment was applied in triplicate to oocytes from each female. After 48 hours of incubation at 21°C, the follicles were examined under a binocular microscope and oocytes without the follicular layer (ovulated) and intact follicles (un-ovulated) were counted for each well.

### 2.3. Egg collection and incubation

Surface out-flow egg collectors (mesh size of 500 μm) were placed to receive eggs from the tanks and were frequently inspected for eggs. When spawning was observed, eggs were transferred into a 10 L bucket. The number of eggs per spawn (fecundity) was estimated by counting the eggs in triplicate subsamples. A sample of eggs (n = 50 – 100) was observed under a microscope and the percentage (%) fertilization for each batch of spawned eggs was determined by calculating the % of eggs that reached the 2- to 16-cell stage. Eggs for incubation were collected after careful agitation of the eggs in the 10 L bucket. The eggs were incubated at a density of 13,230 ± 7,273 eggs L^-1^ in 30 L incubators with the same conditions as the broodstock holding tanks. Each incubator was supplied with an air stone placed down in the centre to maintain the eggs in suspension and prevent accumulation of the eggs at the surface or bottom of the incubator. The number of eggs in each incubator was estimated by mixing the incubator homogenously and taking three 100 mL samples and counting the eggs in each sample. The eggs were left one day to develop and survival rate (percentage of eggs with embryos) was estimated as for percentage fertilization with a sample of eggs (n = 50 – 100) taken from the incubator. The following day, the number of hatched larvae in each incubator was estimated volumetrically as for the eggs. In parallel to the 30L incubators, fertilised eggs were transferred into individual wells filled with sterile seawater in a 96-well cell culture plate (EIA plate) and placed in a refrigerated incubator at 21°C in duplicate for each spawn and revised daily until the last larva died, to estimate hatching rate and larval survival under starvation as in Giménez *et al*. (2006). Percentage of survival was calculated as the number of larvae alive / total hatched larvae.

A preliminary trial was made to examine the larval growth and development. The trial did not focus on survival as the facilities and staff were not available to provide optimal conditions for the larvae. Larval rearing was carried out using mesocosm conditions (Divanach and Kentouri, 2000), with low larval density in a large tank (6 larvae L^-1^, 1500 L tank) under more natural or, at least, less strict conditions than those used in intensive rearing, and using an endogenous bloom of wild marine zooplankton, mostly harpacticoid copepods, together with periodic addition of rotifers and Artemia spp nauplii. The trial was carried out from November 11^th^ to December 18^th^ 2020 using larvae 4 days post hatch (dph) that hatched on November 7^th^, from a spawn obtained with the rLh + rLh spawning treatment. The larvae were stocked in a 1500 L tank at 20°C, 12hL:12hD photoperiod and fed on rotifers for 26 days (4 to 30 dph) followed by newly hatched Artemia nauplii (24 - 39 dph). Phytoplankton (a mixture of *Tetraselmis suecica* and *Isochrysis galbana*) was added every day in order to maintain a green medium, and every two days the rotifer concentration was assessed to maintain a density of 5 rot mL^-1^. Artemia nauplii were added when the larvae reached 4.3 mm TL (20 dph) being the main food for larvae after 5 mm TL as suggested by Hagiwara *et al*. (1992).Ten larvae arbitrarily chosen were sampled at 4, 6, 9, 11, 17, 23, 27, 32, 37 and 39 dph and anaesthetised with MS-222. Standard length was measured using a digital camera connected to an image analyser (AnalySIS, SIS Gmbh, Germany). Photographs were also used to estimate the presence or absence of food in the gut and to examine swim bladder inflation as well as other indicators of larval development.

### 2.4. Histological analysis

Ovarian samples obtained by cannulation and testis portions were preserved in Bouin’s fluid for 24 h and stored in 70% ethanol until processed. The dehydrated tissues were embedded in paraffin and 3 μm sections cut. The testes portions (from the anterior, middle and posterior part) were oriented to obtain horizontal sections. Cut sections were stained with hematoxylin and eosin (Casa Álvarez, Spain) for morphological evaluation. The slides were examined under a light microscope (Leica DMLB, Houston, USA).

Oocytes were classified as previously described by Ramos-Júdez *et al*., (2021). Oocytes were classified as previtellogenic: with primary growth (PG) oocytes, which presented multiple nucleoli situated in the germinal vesicle, or with cortical alveoli (SGca) oocytes that had small oil droplets and granular vesicles in the peripheral ooplasm. The incorporation of yolk globules indicated vitellogenic stages: early secondary growth (SGe), late-secondary growth (SGl) when oocytes reached ≥ 400 μm (Greeley et al., 1987) and full-grown secondary growth oocytes (SGfg) when the vitellogenic growth was completed with the fusion of yolk granules and thickening of vitelline membrane. Oocytes were classified as oocyte maturation stage (OM) when oil droplets were coalescing and the nucleus was positioned to one side of the oocyte, indicating the initiation of the germinal vesicle migration (GVM). Oocytes with disintegrating structure and hypertrophy were in atresia (Valdebenito et al., 2011). Maturation stage of females was determined according to the most developed stage of oocytes present. Additionally, the percentage of oocytes in different stages in the ovaries among weeks was calculated through the identification of ≥ 50 random oocytes per female each week.

To evaluate testes samples, the number of spermatogonia (SPG) type A and B, spermatocytes (SPC), spermatids (SPD) and spermatozoa (SPZ) were scored in 12 seminiferous tubules randomly selected from different areas (anterior, middle and posterior) per sample and the percentage abundance of each germ cell type was determined.

### 2.5. Sperm quality evaluation

Sperm samples from running males were collected, for quality evaluation, on week 10 and week 13 at the end of the experimental period. Samples were collected in a 1 mL syringe avoiding the contamination by urine, faeces and water. Sperm was divided into two aliquots, one was maintained as undiluted sperm and one was diluted 1:10 (1 part sperm + 9 parts diluent) in Marine Freeze® (IMV Technologies, L’Aigle, France) extender (González-López et al., 2020) and both samples were maintained in Eppendorf tubes at 4°C until evaluation. Sperm was activated by pipetting 5 μL of the sperm sample (undiluted or diluted) into an Eppendorf with, depending on the concentration of the sperm, 195, 295, 495 or 995 μL of sea water. Immediately, the Eppendorf was agitated to thoroughly mix, then 2 μL containing activated sperm was pipetted into an ISAS counting chamber (Integrated Sperm Analysis System, Spain), and videos of tracks of the activated spermatozoa were recorded 15 s after activation with the CASA system SCA-VET-01 (Microptic, Barcelona, Spain). Videos were recorded using a digital camera (Basler Ace ACA1300-200UC, Basler AG, Ahrensburg, Germany) connected to an optical phase-contrast microscope (Nikon Eclipse Ci, Tokyo, Japan) with ×10 negative phase contrast objective. The following sperm parameters were determined: (1) sperm concentration (spz mL^-1^), (2) sperm motility (%), (3) rapid progressive sperm (%), and (4) sperm velocity (μm s^-1^): the curvilinear velocity (VCL), straight-line velocity (VSL) and average path velocity (VAP). The CASA program was set to classify motile sperm to have a VCL of > 25 μm s^-1^ and fast progressive sperm to have a straightness (SRT = VSL/VAP x 100) of > 80% and a VCL of > 80 μm s^-1^. All samples were analysed on the day of the collection and 48 h after collection. Samples collected at the end of the experiment (week 13) were also analysed on days 1, 4, 6, 8, 11, 13 and 15 after collection.

### 2.6. Steroid hormone determination

Blood samples were collected and centrifuged at 3,000 rpm at 4 °C for 15 min and the plasma stored at −80 °C until analysis. Plasma levels of 17β-estradiol (E_2_) and 11-ketotestosterone (11-KT) were analysed using commercially available enzyme immunoassays (EIA) (Cayman Chemical Company, USA). Steroids were extracted with methanol that was evaporated and extracts were re-suspended 1:10 (E_2_) or 1:100 (11-KT) in the EIA buffer.

### 2.7. Statistical analysis

Data is expressed as the mean ± standard deviation (SD). A Chi-square test was used to examine the distribution between groups, of fish that had different maturational stages at the start of the experiment and whether fish that ovulated or spawned had a determinate maturity status at the beginning of the experiment. Shapiro-Wilk and Levene tests were used to check the normality of data distribution and homogeneity of variance, respectively. Mann-Whitney U test or Kruskal-Wallis test, followed by Dunn’s pairwise comparison, were used in non-normally distributed data to compare oocyte diameter between treated and control group within a week and between weeks during the experiment, respectively. One-way ANOVA followed by Holm-Sidak *post hoc* test was used to separately examine differences in the independent variables of diameter of full-grown oocytes, percentage of OM, total eggs per females, and fecundity between the three spawning treatments. Variances across the groups were not equal for the GSI data, which was log-transformed and groups compared using the Brown-Forsythe test and Games-Howell *post-hoc* multiple comparisons test. Spawning data from rLh + rLh and rLh + P_4_ spawning treatments (i.e., latency period, fertilization and hatching percentages) was compared using a t-student test. Differences in percentage of OM and oocyte diameter before and after the priming or resolving injections, and percentage ovulation of oocytes incubated *in vitro* were examined by one-way repeated measures (RM) ANOVA with individual females as the subject. Two-way RM ANOVA with pairwise comparison by the Holm-Sidak test was used for comparing E_2_ and 11-KT between weeks and treatment groups. Sperm quality parameters were compared with a one-way and two-way RM ANOVA. In the one-way RM ANOVA, the male was the subject, day of storage the independent variable and sperm quality parameters the dependent variables. In the two-way RM ANOVA, the male was the subject, time of storage (0 or 48h) and sample dilution (undiluted or Marine Freeze®) were the independent variables and sperm quality parameters the dependent variables. Significant differences were detected at a significance level of P < 0.05. Statistical analyses were performed with SigmaPlot version 12.0 (Systat Software Inc., Richmond, CA, USA), with the exception of the Brown-Forsythe test that was conducted with SPSS software version 20.0 (Armonk, NY: IBM Corp).

### 2.8. Ethics statement

The study was conducted in accordance with the European Directive 2010/63/EU of 22nd September on the protection of animals used for scientific purposes; the Spanish Royal Decree 53/2013 of February 1st on the protection of animals used for experimentation or other scientific purposes; the Catalan Law 5/1995 of June 21th, for protection of animals used for experimentation or other scientific purposes and the Catalan Decree 214/1997 of July 30th for the regulation of the use of animals for the experimentation or other scientific purposes. The procedures used were evaluated by IRTA’s Committee of Ethics and Experimental Animal (CEEA) and the Catalan Government and procedures authorized with ID V7MH4802M.

## 3. Results

### 3.1. Induction and completion of vitellogenesis

Induction and completion of vitellogenesis (603 ± 8 μm) from previtellogenic and early vitellogenic females was achieved with 100 % success in the rGth-treated group; as observed for all 12 previtellogenic females and all 9 early vitellogenic females. No females from the control group completed oocyte growth (Fig 4A).

**Figure 4.**
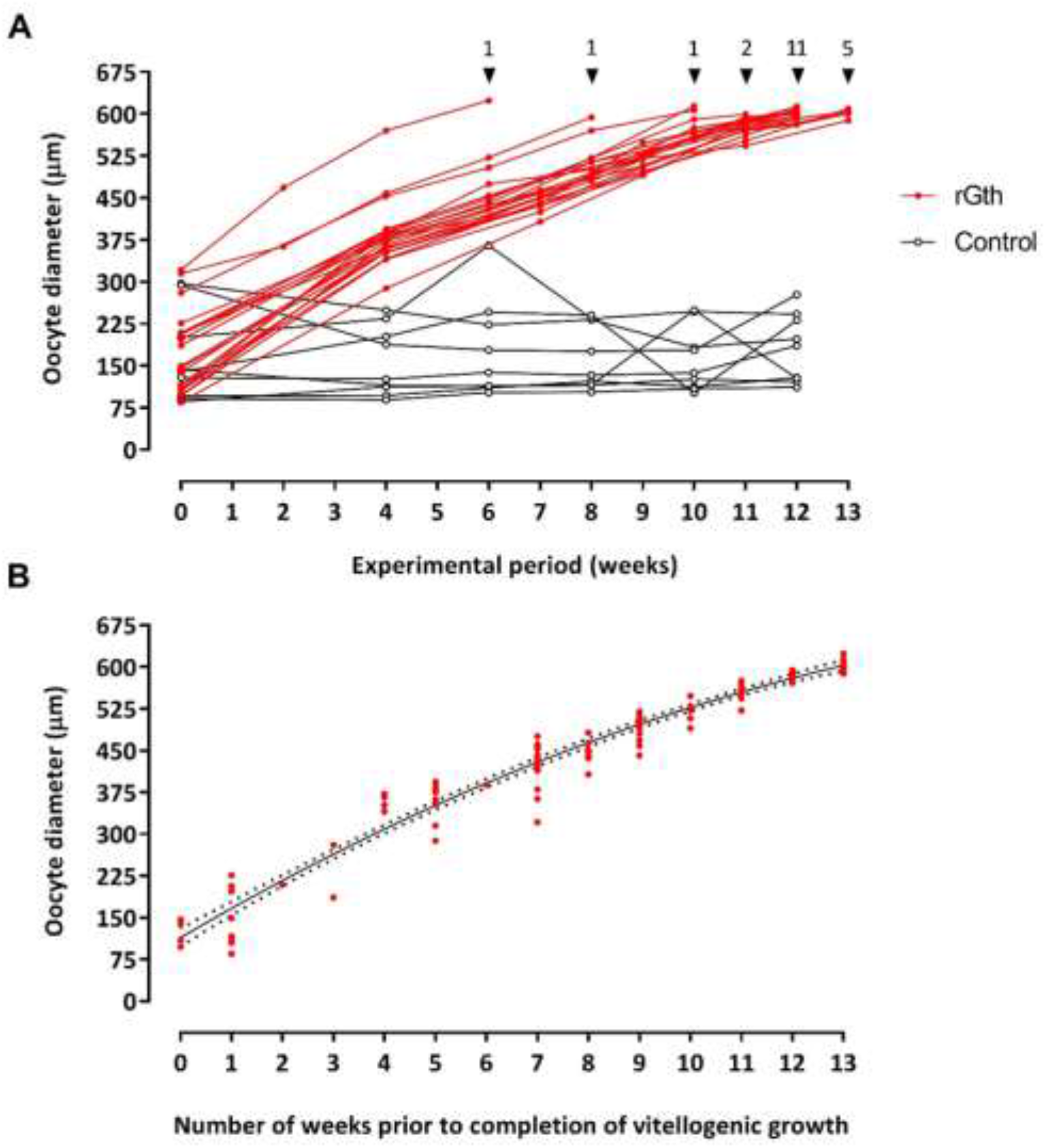
Mean oocyte diameter of the largest most developed oocytes (n = 20 per female) in wet mounts in each female (represented by a line). (A) Females that received the rFsh and rLh treatment (n = 21) and females that received saline injections (control) (n = 9). Triangles show the moment and the number of females that completed vitellogenic growth and were selected for maturation and ovulation induction. (B) Mean diameter of the oocytes of rGth-treated females aligned from the completion of vitellogenic growth, the moment that were selected for maturation and spawning induction. The growth of oocytes in represented by a second order polynomial (quadratic) equation (R^2^ = 0.9625).

At the beginning of the experiment, females in different stages of ovarian development were evenly distributed between the control and treatment groups (χ^2^ = 0.824; gl = 2; P = 0.662). Control and rGths groups had 62 ± 7 % of females in previtellogenesis (from which the 61 ± 7 % and 23 ± 16 % were at primary growth and at cortical alveoli, respectively) and 38 ± 7 % in early vitellogenesis (Fig 5A, 5B) with no significant difference in mean initial diameter of the largest oocytes (control = 164 ± 82 μm, rGth-treated = 172 ± 72 μm). Although some females had started vitellogenic growth, at week 0 the ovaries had principally PG oocytes, some cortical alveoli oocytes (SGca) with very few oocytes in early vitellogenesis and some atresia (Figs. 5C, 5D, 6A).

**Figure 5.**
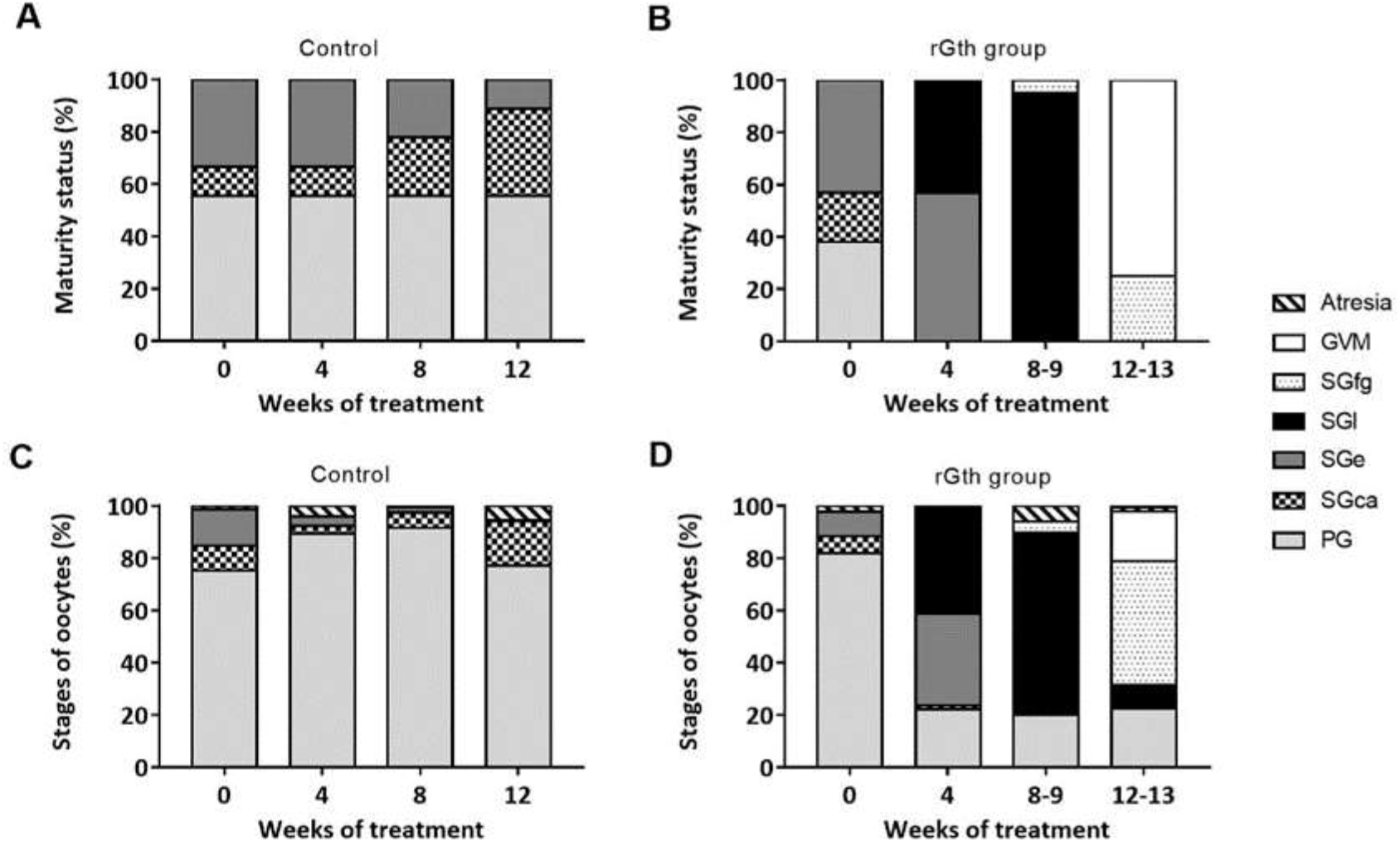
Percentage of females in different maturity stages of gonadal development in control (A) and rGth-treated group (B), and evolution of percentage frequency of oocyte developmental stages observed in control (C) and rGth-treated group (D) in different weeks of the experimental period. The maturity status of females was determined by the most advanced oocyte stage present in the samples. To calculate the percentage of each oocyte stage, a total of 50 random oocytes per female were classified and proportions were estimated. Each bar section represents the mean percentage of oocytes per stage from females for each week. PG, primary growth oocyte; SGca, cortical alveoli step; SGe, early secondary growth; SGl, mid- to late secondary growth oocyte (> 400 μm); SGfg, full-grown secondary-growth oocytes; GVM, initial germinal vesicle migration.

**Figure 6.**
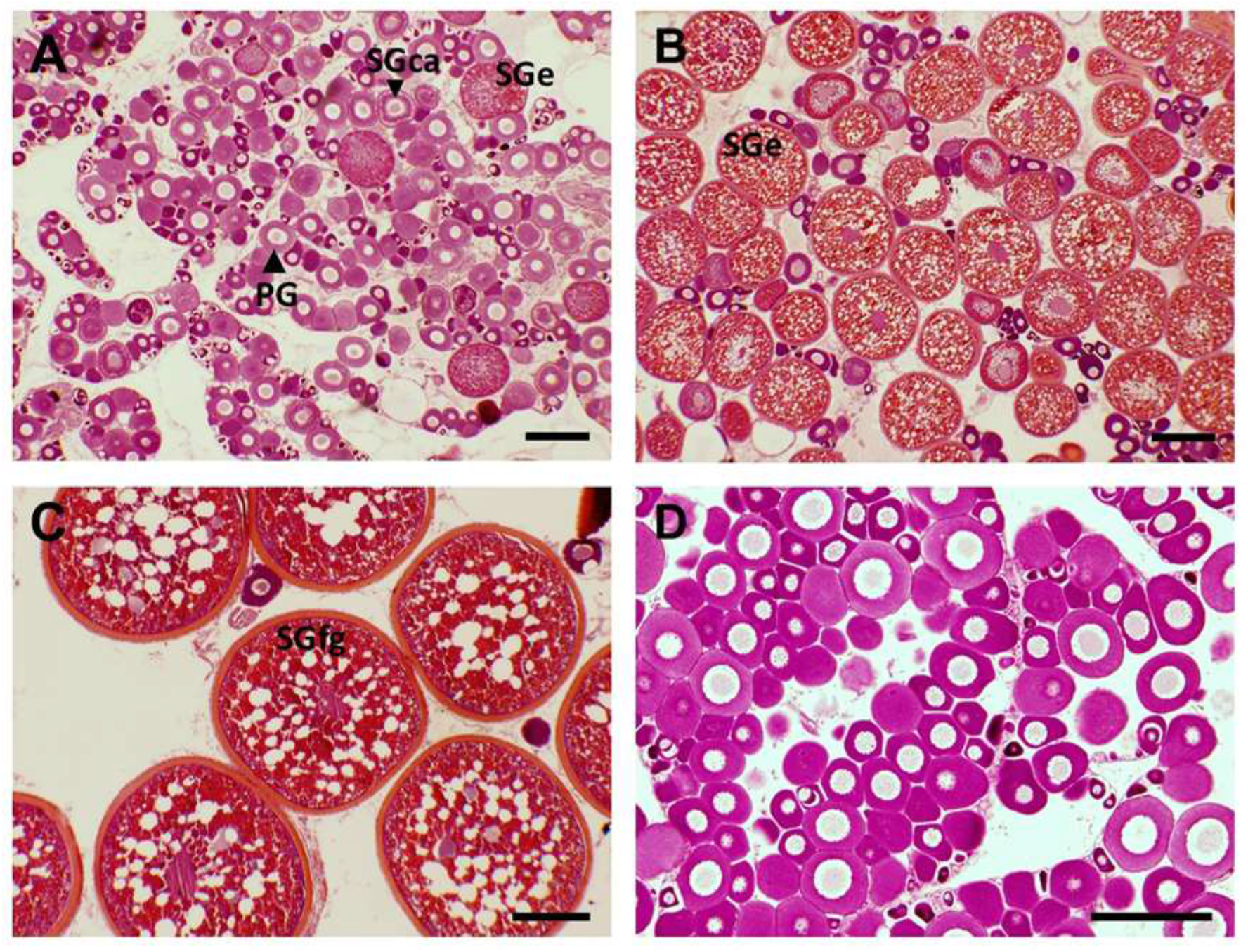
Histological photographs of representative ovarian biopsies from flathead grey mullet females (Mugil cephalus) during the experimental period. (A) A female in early-vitellogenesis (SGe) with mainly primary growth oocytes, some cortical alveoli stage (SGca) and only a small percentage of oocytes in SGe at the beginning of the experiment, (B) a female in SGe after rFsh treatment with a clear clutch of oocytes recruited into vitellogenesis, (C) a rGth-treated female with full-grown oocytes (SGfg), and (D) a control female in previtellogenesis by the end of the experimental period. Scale bar: 200 μm.

Previtellogenic females at PG from the control group did not show further development during the experiment (Fig 5A, 6D). The number of early-vitellogenic females in the control group decreased during the experiment, concomitantly there was an increase in the occurrence of follicular atresia (Fig. 5A, 5C). In addition, no significant differences in oocyte diameter were observed amongst weeks in the control group (Fig 4A). On the other hand, the hormonal treatment induced a significant (P <0.001) growth of oocyte diameter. The rGth-treated group showed a significant weekly increase in oocyte diameter (P < 0.001) (Fig 4A) and by the fourth week of treatment all (100 %) females had progressed to vitellogenesis (Fig 5B). At this point, a clear clutch of vitellogenic oocytes was the most abundant in the ovaries (Fig 5D, 6B). From the sixth week onwards, different females completed oocyte growth (Fig 6C) with the majority completing gonadal development in week 12 and 13. To define the pattern of oocyte growth in the rGth-group, oocytes were aligned from the completion of vitellogenic growth and the growth was represented by a second order polynomial (quadratic) equation y = 113.9 + 53.62 x – 1.234 x^2^ (R^2^ = 0.9625) (Fig 4B). A greater increase in oocyte diameter was observed between weeks (injections) in earlier stages, with ~ 50 μm per week, while approximately three weeks (injections) was required to grow the last 50 μm to complete vitellogenic growth. Once calculated the oocyte volume (V = 4/3 πr^3^), conversely, showed a greater weekly increase (more than 5 times greater) in the later stages of development compared to the early stages.

Concerning E_2_ levels, no significant differences were observed between control and treated group at the beginning of the experiment (0.67 ± 0.71 ng mL^-1^ and 0.68 ± 0.46 ng mL^-1^, respectively), whereas a significant increase (P < 0.001) was observed when the treated group completed vitellogenic growth (1.53 ± 0.70 ng mL^-1^) in comparison with no changes in controls (0.52 ± 0.25 ng mL^-1^) at the end of the experiment.

### 3.2. Induction of spermatogenesis and spermiation

Spermiation was induced in 100% (n = 9) of the males in the rGth-treated group. All males advanced from a sperm index of 0 or 1 at the start of the experiment, indicating no sperm sample could be obtained, to a sperm index of 2 or 3 indicating males had fluent or very fluent sperm. All control males were in sperm index 0, no presence of sperm at the end of the experiment.

At the beginning of the experiment, males with or without presence of sperm were evenly distributed between the control and treatment groups (χ^2^ = 0.156; gl = 1; P = 0.693). Only 20 % (3 out of 15 individuals; 1 from the control group and 2 from the rGths group) presented traces of high viscous milt, but not sufficient to obtain a sample (Fig 7).

**Figure 7.**
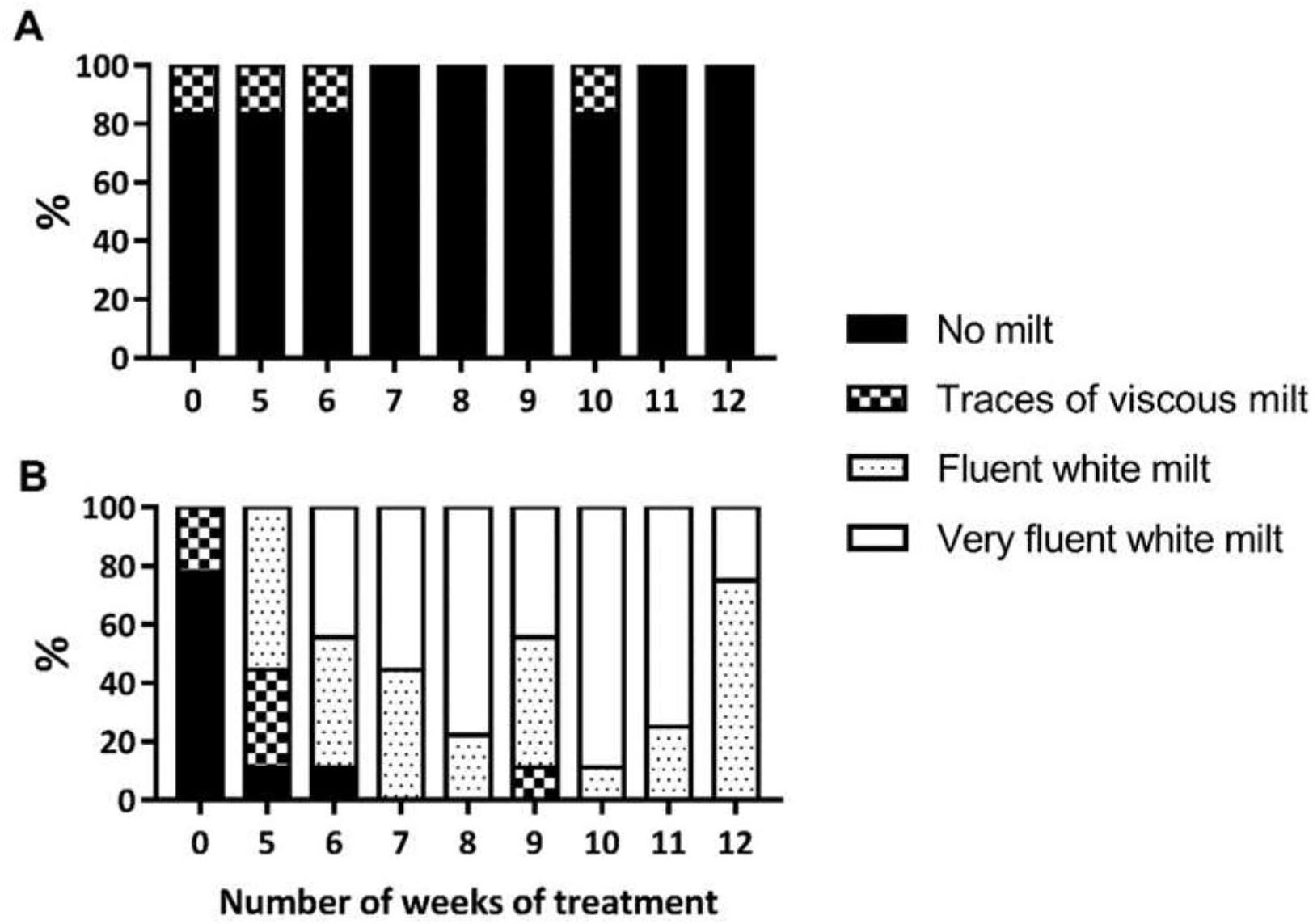
Percentage of no spermiating and different degrees of spermiating males in (A) control (n = 6) and (B) rGth-treated group (n = 9) during the males’ treatment period. Males were classified as no milt (index 0), traces of viscous milt from which no sample could be obtained (index 1), fluent white milt (index 2) and very fluent white milt (index 3). Data from Group 1 (n = 4 rGth-treated males, n = 3 control) from the first application (week 0) to the last checking (week 12) was combined with data from Group 2 (n = 5 rGth-treated males, n = 3 control) from week 0 to 10 in order to present the results.

The histological evaluation of the testes from two fish in which no milt was obtained after abdominal pressure at the beginning of the experiment, showed that one male presented only SPG within the seminiferous tubules, while the other male presented many SPG and some SPC, SPD and SPZ, but many tubules did not have the central lumen formed and those that had, presented very few spermatozoa (Fig 8A, B). During the experimental period, males in the control group did not produce fluent milt and only the same one male out of six (17%) produced a small drop of viscous milt within different weeks.

**Figure 8.**
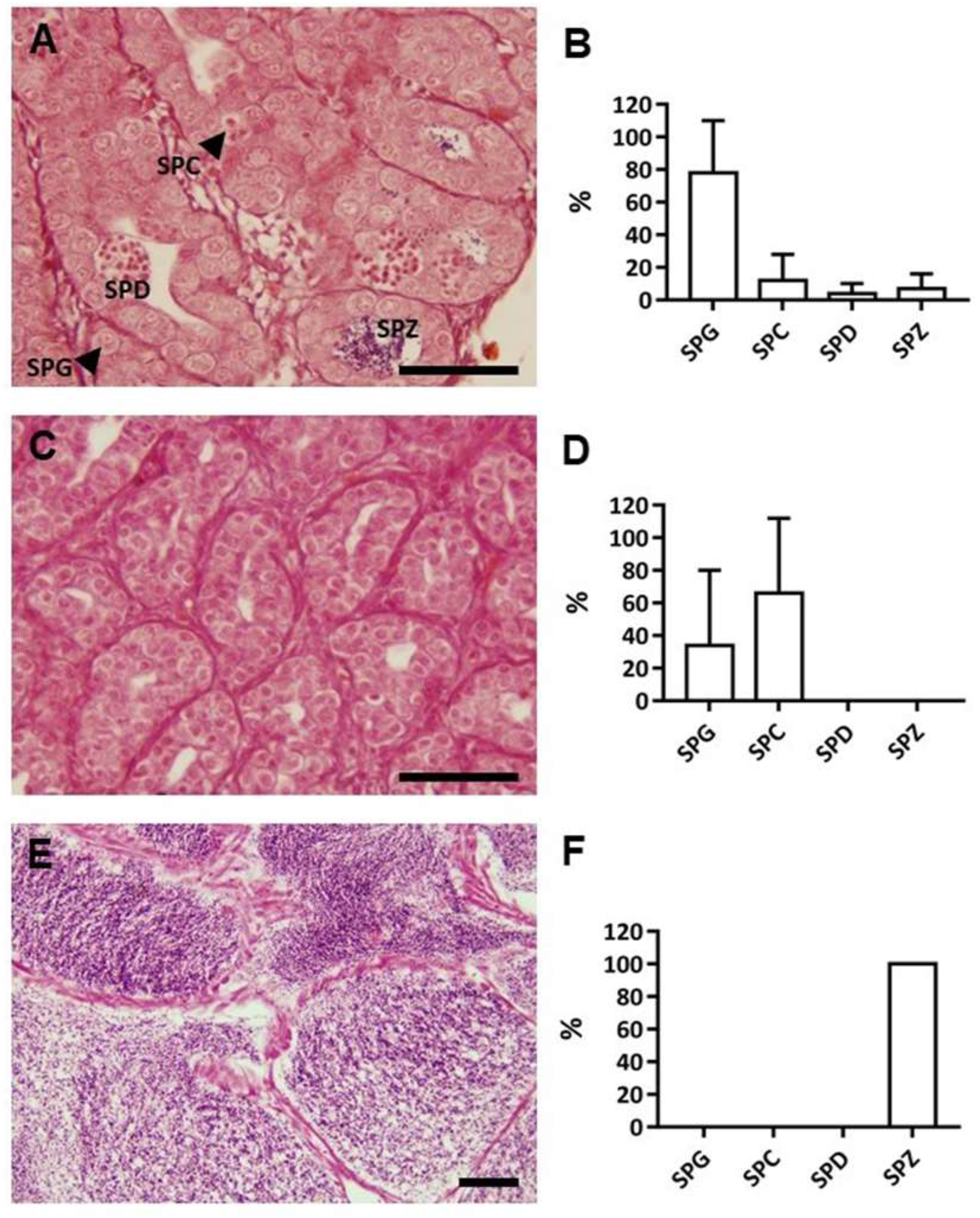
Histological sections of testis and the percentages of the spermatogenesis developmental stages at the beginning of the study (A and B, n = 2) and at the end of the experimental period in control group (C and D, n = 2) and rGth-treated group (E and F, n = 2). SPG, Spermatogonia; SPC, Spermatocyte; SPD, Spermatid; SPZ, Spermatozoa. Scale bars = 50 μm.

At the end of the experiment, the testis from two control males —the male that in previous weeks presented viscous milt and one male that never had milt— contained tubules with a higher number of SPC than the initial situation, but there was no presence of SPD or SPZ (Fig 8C, D). In contrast, running males with fluent white milt were observed in the rGth-group. After five weeks of treatment (three weeks of rFsh treatment and two weeks of combined rFsh and rLh treatment) eight out of nine (89 %) males presented milt; either viscous traces (33 %) or fluent milt (56 %) (Fig 7). With the application of rLh, the number of males with fluent milt increased. By seven weeks, 100% of the males were spermiating with fluent milt and males maintained fluent milt until the end of experiment (week 12), with the exception of week 9 when one male produced viscous milt. However, after the application of rFsh in Group 1 and continuing the rLh treatment in Group 2, fluent milt was produced again in the following week. In clear contrast to the control males, in the histology from two rGth treated males at the end of the experiment, the sperm ducts of the rGth-group were completely filled with spermatozoa (Fig 8E, F) and sperm volumes ranged from 0.25 to 2.89 mL. In addition, GSI values reflected the growth of the testis with rGths treated males having significantly higher (P = 0.026) GSI compared to control group at the end of the experiment. Males before the hormonal treatment (n = 2) and control males at the end of the treatment (n = 2) showed thin undeveloped testes with GSI of 0.10 ± 0.05 % and 0.06 ± 0.01 %, respectively, while rGth-treated males at the end of the experiment (n = 2) presented well developed white testes with 5.35 ± 1.25 % GSI. Taken together, males with sperm index 0 and 1 had germ cells predominantly at stages of SPG and SPC in undeveloped testes (GSI ≲ 0.1) and the rGth treatments induced testes growth (GSI 5.35 ± 1.25 %) and development of germ cells to complete spermatogenesis and provide testes full of spermatozoa and fluent spermiation (index 2 and 3).

In regard of steroid changes, rGths treatment had a significant effect in the increase of 11-KT levels (P < 0.001) compared to the control group (Fig 9) that was maintained without significant changes. In the rGths-group, increasing values of 11-KT were obtained within the course of the experiment coinciding with the availability of spermiating males, with the maximum at eight weeks and a decrease afterwards.

**Figure 9.**
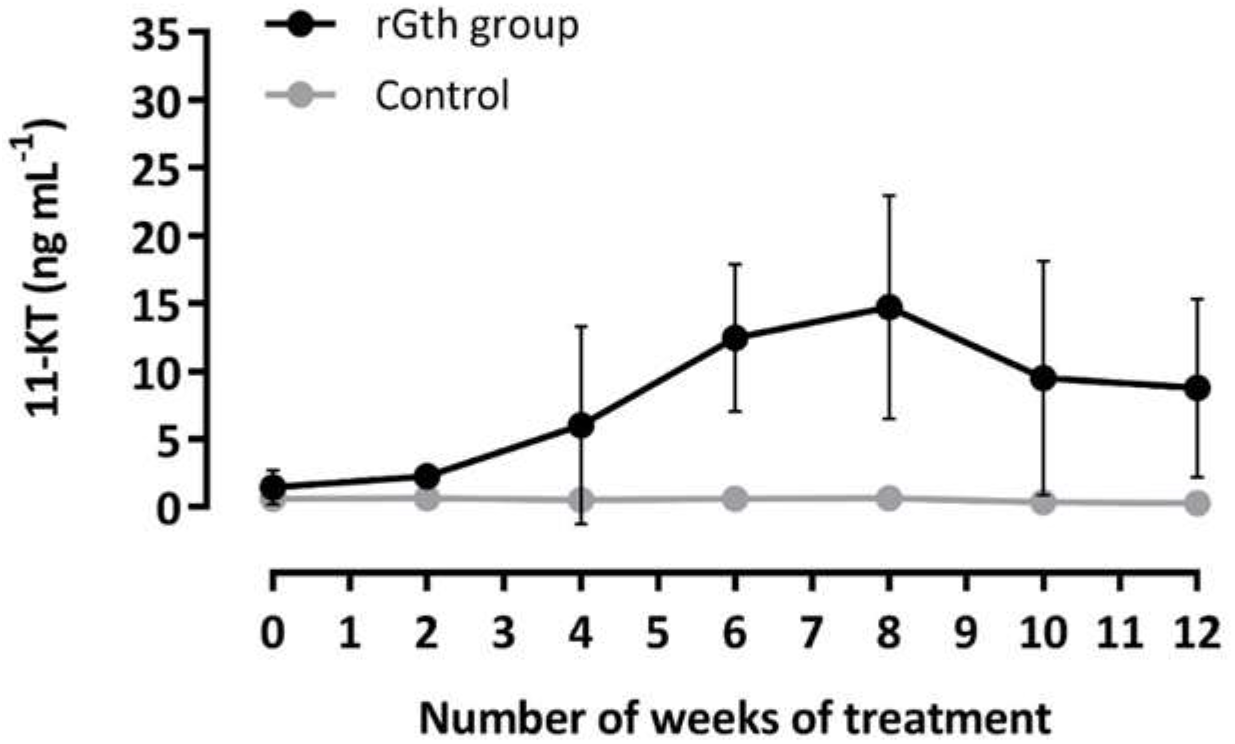
Effect of rFsh and rLh treatment (rGth group) and saline (control) on 11-ketotestosterone (11-KT) levels (mean ± SD) in the flathead grey mullet (*Mugil cephalus*). Data from Group 1 (n = 4 rGth-treated males, n = 3 control) from the first application (week 0) to the last checking (week 12) was combined with data from Group 2 (n = 5 rGth-treated males, n = 3 control) from week 0 to 10 in order to present the results. Two-way RM ANOVA showed a significant effect of the rGths treatment on the production of 11-KT (P < 0.001).

Regarding sperm quality, on the day of collection (0h), the sperm diluted 1:10 in Marine freeze had the following mean characteristics: motility of 58 ± 22 %, head size of 13 ± 5 μm^2^, 107 ± 24 μm s^-1^ VCL, 92 ± 29 μm s^-1^ VAP, 70 ± 31 μm s^-1^ VSL, 67 ± 12 % STR, 57 ± 16 % LIN, and 79 ± 11% WOB. Whilst 20 ± 16 % of the motile spermatozoan were fast progressive that had velocity of 149 ± 17 μm s^-1^ VCL, 140 ± 20 μm s^-1^ VAP, 129 ± 21 μm s^-1^ VSL, 92 ± 2 % STR, 86 ± 7 % LIN, and 93 ± 6% WOB. The variation amongst the nine males was wide with % sperm motility ranging from 19 to 89%. The mean concentration was 7.59 x 10^10^ spermatozoa mL^-1^ or 15.56 x 10^10^ spermatozoa kg^-1^. The sperm diluted in Marine Freeze® had significantly higher motility than undiluted sperm on the day of collection (0 h) and 48 h after collection (Fig 10). The motility of undiluted sperm decreased significantly from day 0 to 48 h after collection, compared to diluted sperm that maintained similar motility. The sperm collected and diluted in Marine Freeze® at the end of the experiment (week 13) maintained similar motility for 6 days of storage at 4°C (tested days 0, 1, 2, 4 and 6) with variation from 41.1 ± 20.0% to 70.2 ± 17.3% before decreasing significantly from day 4 (70.2 ± 17.3%) to day 8 (11.7 ± 5.1%), whilst day 6 was intermediate (41.1 ± 20.0%).

**Figure 10.**
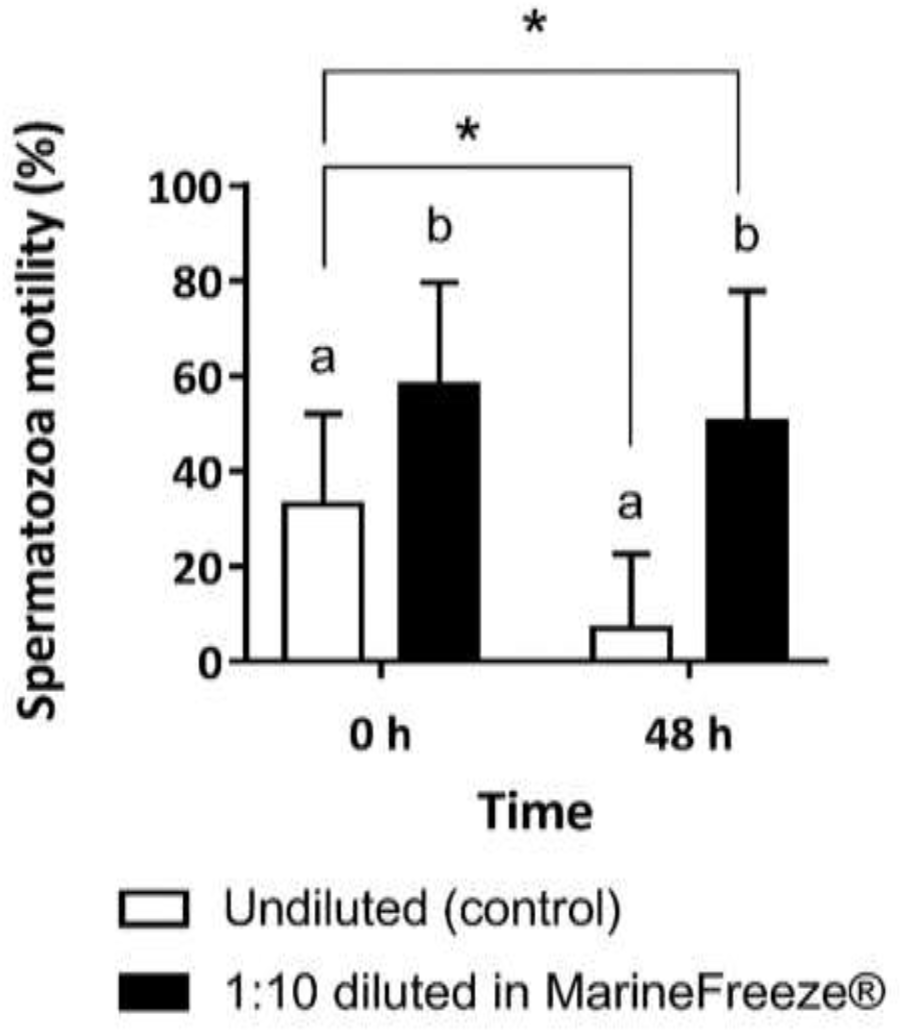
Percentage motility of sperm samples collected from rGth-treated males at the end of the experimental period (week 10 and 13). A two-way RM ANOVA was performed followed by the Holm-Sidak *post hoc* test with males as subjects, time of storage (0 or 48h) and sample dilution (undiluted or 1:10 diluted in Marine Freeze®) as the independent variables, and percentage motility as the dependent variables. Different letters indicate significant differences between undiluted and diluted samples within the same time of evaluation, while asterisks show significant differences between evaluation time.

### 3.3. Oocyte final maturation and spawning induction

Females of flathead grey mullet that had completed vitellogenic growth —available only from the rGth-group— were selected for spawning induction following three different treatments. Gonad samples were examined before the priming injections and after 24:05 ± 0:40 h, just prior to the resolving doses. Ovaries from chosen females were composed by SGfg with a mean size of 603 ± 8 μm and the 76 % (16 of 21) of the females presented 17 ± 23 % of oocytes that had already initiated OM with the migration of germinal vesicle (Fig 5D); the nucleus had moved off-center and there was a small degree of fusion of yolk granules and lipid droplets, but with no single large yolk mass. No significant differences in oocyte diameter or the percentage of OM were found between the females that received different spawning treatments. After the priming injections (24 h), 100% of the females that received rLh had entered into OM with most of the oocytes (96 ± 9 %) in late GVM with yolk coalescence (Fig 11). Meanwhile, those that received P_4_ did not show a significant increase in GVM percentage (35 ± 35 %) respect to initial stage (12 ± 5 %) thereby showing that most of the oocytes were retained at the secondary growth stage.

**Figure 11.**
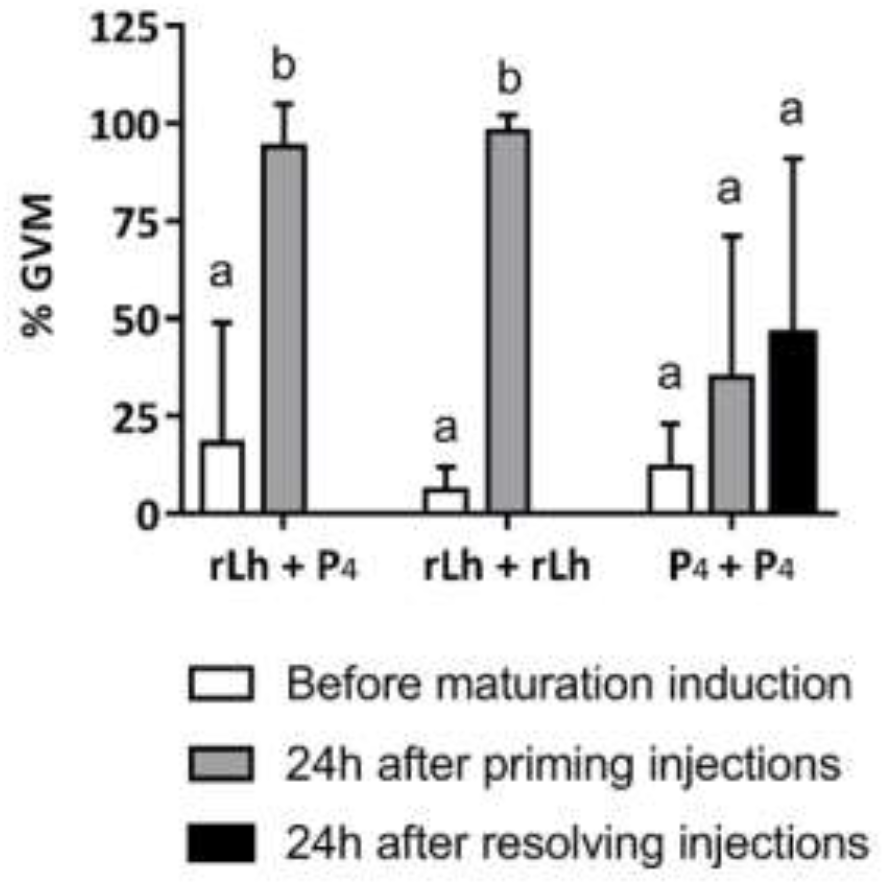
Percentage of oocytes at maturation stage (OM) with germinal vesicle migration (GVM) before OM induction, 24 h after the priming injections and 24 h after the resolving injections. Treatments applied to induce OM and spawning were (i) priming 30 μg kg^-1^ rLh and 40 mg kg^-1^ resolving Progesterone (P_4_), (ii) 30 μg kg^-1^ rLh as priming and resolving injections or (iii) 40 mg kg^-1^ P_4_ as priming and resolving injections given 24:05 ± 0:40 h apart. Different letters indicate significant differences at different timing of the inductions within a same treatment following One-way RM ANOVA.

The proportions of females that ovulated or spawned did not vary between the initial maturity status of females at the beginning of the experiment (χ^2^ = 1.150; gl = 2; P = 0.563 0.001; χ^2^ = 2.149; gl = 2; P = 0.342). In the rLh + P_4_ group, 100 % of the females ovulated and eight out of nine females (89 %) spawned in the tank 16:36 ± 1:56 after the resolving injection. In the rLh + rLh group, 100 % of the females (n = 6) ovulated and spawned 16:51 ± 2:22 h after the resolving injection. Meanwhile, three out of six fish (50%) under P_4_ + P_4_ treatment ovulated but only one (17%) actually liberated some eggs (177,667 eggs) 24:30 h from the administration of the resolving dose. The average number of eggs spawned and female fecundity for each hormone treatment were similar (Table 1). Those females that did not ovulate in the P_4_ treatment group, did not show a significant increase in OM after the second P_4_ (Fig 11). The eggs from the females that ovulated and did not spawn were stripped by applying gentle abdominal pressure to liberate all ovulated eggs from the female. The stripped eggs did not have the appearance of viable eggs and formed a dense globular mass with almost no ovarian fluid.

The *in vitro* incubation of oocytes that had initiated OM confirmed the *in vivo* ovulation and spawning. The highest percentages of ovulation (> 50%) were obtained from oocytes treated with P_4_ (4000, 1000, 500 and 50 ng kg^-1^) or 100 ng kg^-1^ of rLh (Fig 12). All oocytes treated with P_4_ and oocytes treated with 400, 200 and 100 ng kg^-1^ of rLh had significantly (P < 0.05) higher percentages (> 34%) of ovulation than untreated oocytes (control) and oocytes treated with rFsh or 10 ng kg^-1^ of rLh (< 8%). Oocytes treated with 50 ng kg^-1^ of rLh had a percentage of ovulation (21.3 ± 18.5 %) that was intermediate between the highest and lowest ovulation groups.

**Figure 12.**
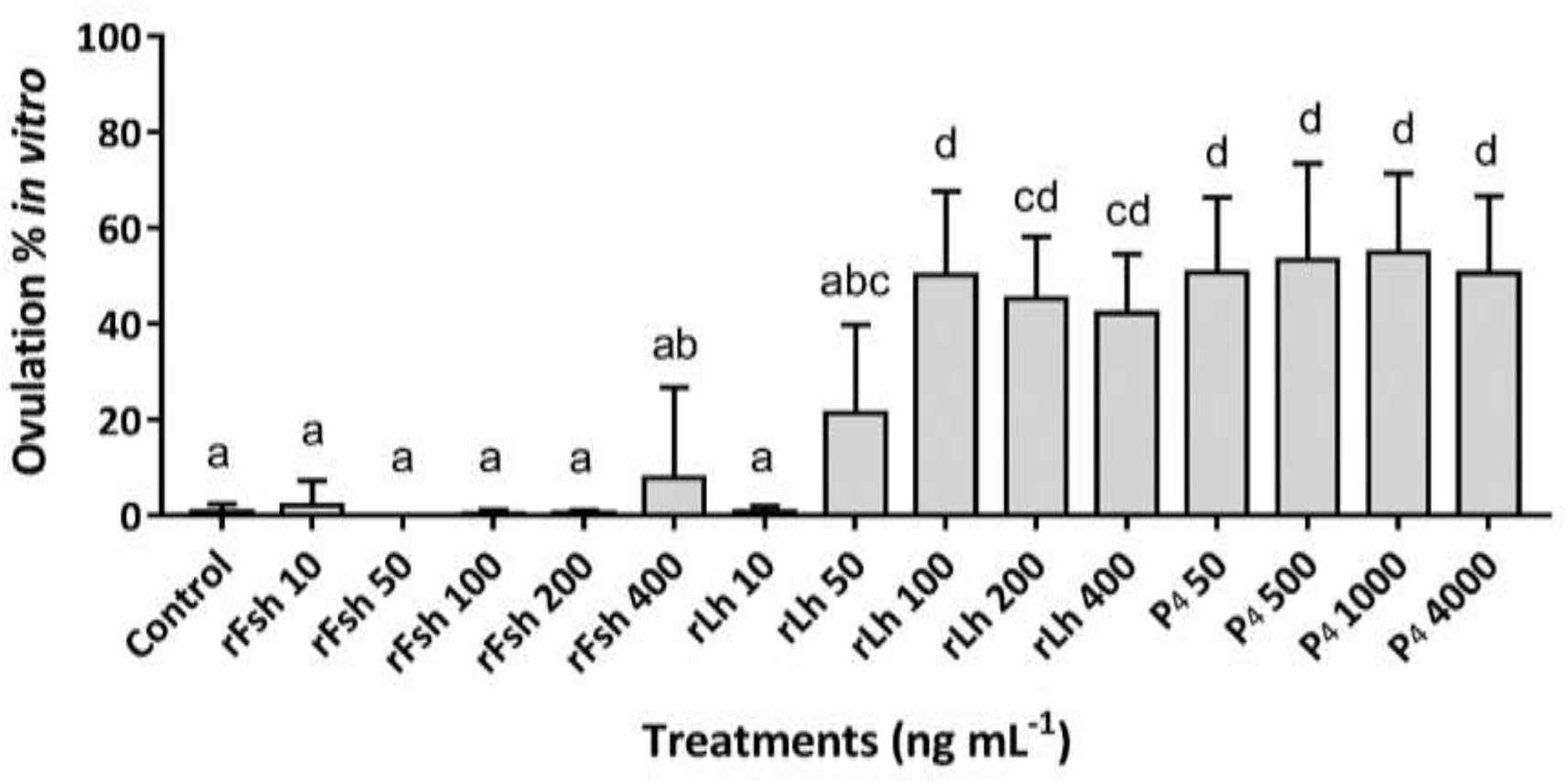
*In vitro* percentage of ovulation (mean ± SD) of oocytes in OM with different effectors, doses and combinations of efectors and doses. Statistical differences between treatments were examined by a one-way repeated measures ANOVA with individual females as subjects (3 replicates per individual, 6 individuals n = 18 wells per hormone concentration).

### 3.4. Egg and larval quality

No significant differences were found in latency, egg production and quality parameters; fecundity, percentages of fertilization, embryo survival or hatching, in those females that received rLh as priming dose and either rLh or P_4_ as resolving injections (Table 1). There were also no differences between females that were at different maturity stage at the beginning of the experiment. Mean total fecundity was 1,738,798 ± 868,950 eggs per female (relative fecundity was 1,245,600 ± 552,117 eggs kg^-1^ bw) with 54 ± 21 % fertilization. On the contrary, spawned eggs from one female that received P_4_ + P_4_, were not fertilised (0% fertilization).

No differences were observed in egg and larval quality between rLh + rLh and rLh + P_4_ groups. The embryo survival and hatching percentages were 64 ± 22 % and 57 ± 24 %, respectively. Hatching in EIA 96-well plates was 96 ± 3 %. Eggs with an embryo measured 0.83 ± 0.02 mm in diameter (Fig 13A) and larvae length at hatching was 1.86 ± 0.14 mm total length (TL). Larval survival rate in the 96-well plates at 21 °C was 85 ± 18 % at 2 dhp, 67 ± 18 % at 5 dhp, 55 ± 17 % at 9 dph and decreased until 8 ± 11 % on 12 dph. By 14 dph, all larvae were dead.

**Figure 13.**
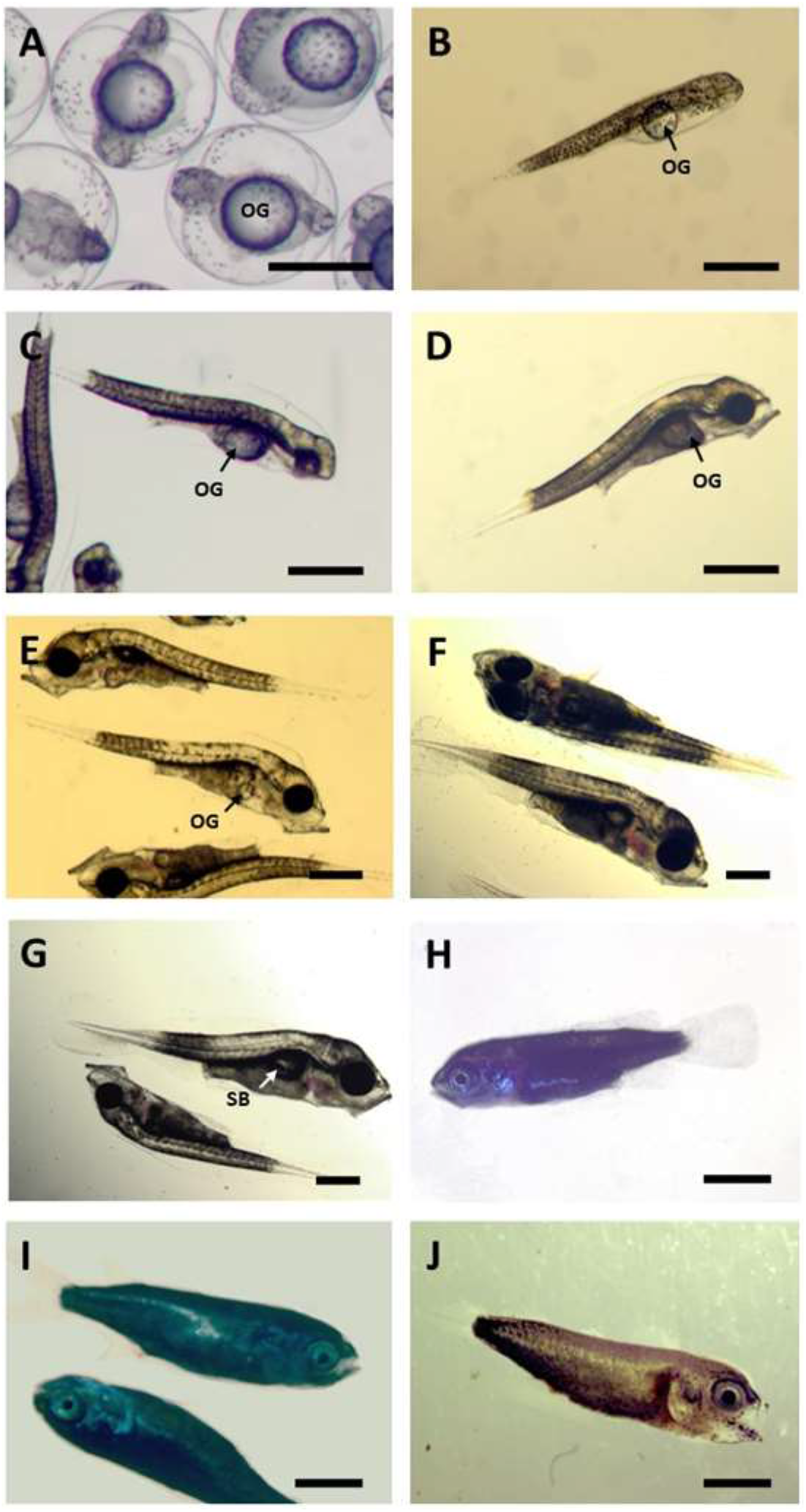
*Mugil cephalus* embryos, larvae, post-larval and juvenile stages. (A) Embryo with head region formed and dark pigment covering almost all the body and the oil globule (OG), (B) newly hatched larvae, (C) larvae at 2 dph with eyes already pigmented, (D) larvae at 3 dph with the mouth parts completely formed and functional, (E) larvae at 9 dph with the oil globule still present, (F) larvae at 19 dph with the oil globule completely absorbed and (G) with a hyperinflated swim bladder (SB), (H) post-larvae at 32 dph, (I) post-larvae at 37 dph, (I) juvenile at 39 dph. Scale bar: 500 μm in A, B, C, D, E, F and G, 1 mm in H, I, and J.

Larvae used for the preliminary larval culture trial measured 2.1 ± 0.05 mm TL at hatching showing a homogeneous yolk mass with a round oil droplet at the posterior part of the yolk sac (Fig 13B). The eyes started to be pigmented between days 1 and 2 dph. Upper and lower jaws were well-developed and the mouth was open from 3 dph (2.93 ± 0.08 mm TL, Figs 13C, 14) when the larvae had consumed most of the yolk reserves and only the oil globule remained (Fig 13D). The pre-flexion stage lasted from 4 to 19 dph with yolk and oil droplet completely absorbed (11 dph). Swim bladder formation started around day 4 - 6 (2.9 - 3.11 mm TL, Fig 14) visible ventrally beneath the notochord. The swimming activity of the larvae increased during the formation and enlargement of the swim bladder although several larvae either swam at the bottom of the tank, or remained floating in the water surface without moving and/or feeding on rotifers due to a hyperinflation of the swim bladder (Fig 13G). Most of these larvae died and, in order to reduce this mortality, light intensity was reduced to 500 lux from 9 dph until the end of the rearing trial. The tail flexion was completed when the larvae reached 3.7 mm length whereas the post-flexion stage was extended during several days (20 - 30 dph, 4.3 - 5.3 mm TL, Fig 14) until the caudal fin and fork were completed. At the end of the trial (day 39) all the fish looked like adult individuals and were considered juveniles (Fig 13J). Development of *Mugil cephalus* larvae and post-larval stages to juvenile are shown in Figs 13 and 14.

## 4. Discussion

Flathead grey mullet (*Mugil cephalus*) held in intensive captive conditions experience reproductive dysfunctions early in the maturation process and both males and females needed to be assisted to induce vitellogenesis, oocyte maturation, ovulation, enhance spermatogenesis, spermiation and spawning. The present study shows that rFsh and rLh can be used as a reliable method to induce and complete oogenesis from previtellogenesis, produce milt and induce spontaneous voluntary tank spawning, in 100% of experimental fish to provide viable, good quality eggs and larvae.

Observing the ovarian developmental stage at the start of the experiment, a range of dysfunctions were found in females; fish were arrested at stages that ranged from previtellogenesis as in Ramos-Júdez *et al*. (2021) through to early vitellogenesis as reported in other Mediterranean areas (Aizen et al., 2005). Nevertheless, in the present study, the ovaries of the early-vitellogenic females at the beginning of the experiment had few vitellogenic oocytes that were not distributed homogeneously and actually previtellogenic oocytes were the most abundant stage of development in all females. The females that received the rFsh and rLh treatment, as previously described by Ramos-Júdez *et al*. (2021), showed a uniform clutch of abundant oocytes recruited into vitellogenesis and followed the typical ovarian development in this species. *Mugil cephalus* has group synchronous ovary development with one clutch of oocytes maturing annually following a single spawning episode (Kumar et al., 2015). The rGth hormonal administration was 100% successful in inducing vitellogenesis from previtellogenesis (12 of 12 females) and early vitellogenesis (9 of 9 females) to complete vitellogenic growth. In comparison, control females that were at early vitellogenesis remained without advances in gonad development during the experiment and presented atresia, which may indicate a lack of stimulation of the vitellogenic oocytes to develop further (Lubzens et al., 2010). Previtellogenic control females did not show further development. This observation is in clear contrast with Aizen *et al*. (2005), in which up to 20% of control females developed mature oocytes without hormonal stimulation. The completion of vitellogenesis in rGth-treated females was accompanied by an increase in plasma E_2_ levels, that was not observed in controls, showing the gonadotropic stimulation of the ovary by rGths. Ramos-Júdez *et al*. (2021), using the same rGths, obtained an almost identical result with an increase in plasma E_2_ levels and eight out of nine (89%) of treated females completing oogenesis.

The rGth treatment in males induced and enhanced spermatogenesis and spermiation as 100% of treated males produced fluent milt for 8 weeks or more. In comparison, control group males remained with no fluent sperm during the experiment. The histological examination of males at different stages of spermiation showed that initial or control captive males that presented no sperm or a little drop of viscous sperm had undeveloped testes with a low GSI (≲ 0.1 %), predominantly SPG and SPC and few or no SPD or SPZ. In comparison, the histological examination showed rGth-treated males had well developed testes with a significantly higher GSI (5.35 ± 1.25 %), that was 50 times higher than control males, and testes were filled entirely with SPZ and no SPG or SPC. Although there was significant growth, some caution is perhaps required in extrapolating the histology from two fish to the other individuals in the control and rGth groups. Few males were available and the n for histology was reduced to ensure sufficient males for rGth treatment and spawning. All the evidence indicates that the rGths have induced spermatogenesis of large numbers of germ cells from early stages (SPG and SPC) through to SPZ. Clearly more work is required for confirmation of this rGth action and a specific study on the direct effect of each rGth on testes development and in the pattern of administration has to be performed to consolidate conclusions. However, other studies, as suggested in the present study, have successfully induced spermatogenesis with rGths, i.e. in sexually immature European eel (Peñaranda *et al*. 2018), immature Japanese eel (Hayakawa et al., 2008; Kamei et al., 2006; Kobayashi et al., 2010) and mature Senegalese sole (Chauvigné et al., 2018, 2017). Peñaranda et al. (2018), represents a specific study that tested different combinations in the application of homologous rFsh and rLh (produced by Rara Avis Biotec, S. L. as in the present study) in immature European eel that lead to different testis development including different degrees of spermiation. The biological effects of rGths on males were also evaluated through plasma 11-KT levels, which is the major androgen responsible for testicular development (Aizen et al., 2005; Chauvigné et al., 2012; Mañanós et al., 2009; Schulz et al., 2010). The rGth treatment significantly increased the levels of 11-KT in the plasma of treated males compared to control males. The concentration of plasma 11-KT in treated males increased gradually in response to rFsh and rLh application, as maturity progressed with the presence of sperm and the increased fluidity of milt obtained in all treated males from 5 weeks onwards. The levels of 11-KT measured, from 0.1 to 28 ng mL^-1^, were in the range of levels previously measured in flathead grey mullet males treated with 17α-methyltestosterone implants to enhance spermatogenesis and spermiation (Aizen et al., 2005).

Higher quantities of fluent milt were obtained (0.25 to 2.89 mL) in the present study compared to Ramos-Júdez *et al*. (2021) (max ~ 0.25 mL). The increased sperm production may be due to the different pattern and dosage of rGths administration, which included a longer period and higher doses for rLh. The quality of the sperm was variable amongst the nine rGth-treated males with adequate mean sperm quality parameters of motility and velocity. The volume of sperm obtained was amongst the highest reported for wild mature *Mugil cephalus*, such as 0.1 to 2 mL (Ramachandran and Natesan, 2016) and the concentration was two powers to ten (10^10^) higher than previously reported (10^8^) (Ramachandran and Natesan, 2016) indicating that a higher degree of dilution during spermiation, which is attributed to the action of Lh (Schulz and Miura, 2002), would be desirable. The extender Marine Freeze® was effective to maintain sperm quality during 6 days of cold 4°C storage, which also indicated the sperm was of good quality.

From an applied point of view, male development was synchronised with the female’s completion of vitellogenesis making them available for the spawning events. Selected males had fluid sperm and were administered either 12 or 24 μg kg^-1^ rLh to stimulate spermiation and reproductive behaviour. Regarding females, three treatments were applied to induce oocyte maturation, ovulation and spawning. The application of 30 μg kg^-1^ rLh as priming injection induced oocyte maturation with the migration of the germinal vesicle in all 100% of females when revised before the application of the resolving dose (24h after the priming dose). The following resolving injections of 40 mg kg^-1^ P_4_ or 30 μg kg^-1^ rLh induced 100 % of the fish to ovulate, and 89 % (P_4_ resolving) or 100 % (rLh resolving) of fish to tank spawn, with a mean percentage fertilization of 54 ± 21 % indicating the success in the application of both treatments. The *in vivo* success of the resolving doses was confirmed by the success of both P_4_ and rLh to induce *in vitro* ovulation after the priming dose of rLh. The *in vitro* test indicated the importance of the selection of a correct rLh dose for the induction of ovulation, as the response of oocytes was different depending on the rLh concentration in the media. Otherwise, low P_4_ doses were as effective for inducing ovulation as high doses which suggests that a refinement in the P_4_ dose applied *in vivo* would be possible. On the contrary to females that received rLh as a priming injection, females that received priming P_4_ had not initiated OM 24 h after the priming dose. The P_4_ resolving dose had no effect on 50 % of females, whilst the other 50 % of the females ovulated, and from this, just one female (17%) spawned eggs that had no fertilization. Although P_4_ as priming and resolving treatment did induce ovulation, all the maturation and ovulation process was concentrated in less than 24 h after the resolving dose was applied. In comparison, females that received rLh as priming completed OM and ovulation during 40:45 ± 2 h initiating OM after the priming dose and completing OM, ovulation and spawning after the resolving dose. It is possible that ovulation was induced by the double P_4_ even though OM had not been induced, indicating the importance in this process of Lh or Lh-induced factors as previously indicated by Ramos-Júdez *et al*. (2021). In that study, only the females that received the highest priming rLh dose (30 μg kg^-1^) proceeded to OM compared to females that received a lower priming rLh dose (15 μg kg^-1^), which did not develop to OM. Taken together, the action of the hormones appear to confirm described roles (Lubzens *et al*., 2010; Mañanós *et al*., 2009) as high doses of rLh were required to induce oocyte maturation and either rLh or P_4_ were needed after the priming rLh dose to induce ovulation and spawning.

The two hormone treatments, rLh + rLh and rLh + P_4_, induced spontaneous tank spawning of large numbers of fertilised eggs. Therefore, the rGth treatment did not only induce gametogenesis development, but also the reproductive behaviour of both sexes to achieve a successful courtship that led to spawning and successful fertilisation of liberated gametes. The presence of good quality males in the tank with fluent milt may also have been a decisive factor for spawning success. Besides, a proper male to female sex ratio could also have been important. In the present study, male to female ratios of 2:1 or 3:1 were placed together and typical mating behaviour —swimming close to the female, pushing the abdomen with their head and body, ceasing to swim momentarily (Whitfield et al., 2012)— was observed by males when females had swollen bellies. In addition, the data from the present study indicate that high-quality eggs up to 80% fertilization can be obtained through the induction of oogenesis by rFsh and rLh in previtellogenic females. Retrieving good quality floating eggs from females that spawned with treated males contrasts notably with the report by Ramos-Júdez *et al*. (2021) in which no spontaneous spawning was observed after priming rLh and resolving P_4_ spawning treatment. Ramos-Júdez *et al*. (2021) did not observe spawning behaviour in rGth treated fish and, therefore, gametes were stripped and artificially fertilised to obtain 0.4 % percentage fertilisation. Therefore, the present study would suggest that other factors such as a delay in the stripping of the eggs coinciding with the process of egg overripening as stated in Ramos-Júdez *et al*. (2021) may have resulted in the low fertilization percentages in that study rather than the application of the rGth treatment *per se*.

Comparisons of spawning success (number of fish that spawned from total injected), fertilization rates and fecundities in other studies with flathead grey mullets are limited owing to differences in methodology and initial gonadal development stage of individuals. A few studies have attempted to enhance vitellogenesis, such as administration of 5 mg kg^-1^ Domperidone (Dom) or in combination with implants of 10 μg kg^-1^ gonadotropin-releasing hormone agonist (GnRHa), in which lower rates of fully mature females were obtained (50% - 85%) (Aizen et al., 2005). Many other studies worked with fully mature females and applied different treatments to induce oocyte maturation and spawning. For example, Aizen *et al*. (2005) applied GnRHa (10 μg kg^-1^ priming, 20 μg kg^-1^ resolving) combined with Metoclopramide (15 mg kg^-1^ priming and resolving), El-Gharabawy and Assem (2006) injected 20 to 70 mg kg^-1^ carp pituitary extract or 10,000 IU fish^-1^ hCG as priming and one or two resolving injections of 100 - 200 μg kg^-1^ GnRHa, while Besbes *et al*. (2020) treated with a priming dose of 10,000 IU kg^-1^ hCG and resolving of 10,000 IU kg^-1^ hCG and 200 μg kg^-1^. Those treatments respectively resulted in: (i) 85% spawning success with low (<40%) to high (< 90%) fertilization percentages and fecundities of 1,649,000 eggs kg^-1^ bw, (ii) 40 % spawning success with 75 to 80 % fertilization and fecundities of 1,395,000 eggs kg^-1^ bw, and (iii) 100 % success with 63 % fertilization but low fecundities of 418,945 eggs kg^-1^ bw. In general, these studies showed a highly variable spawning success and / or variable fertilization percentages whilst the present study presents a reliable spawning success, from 85 to 100% depending on the spawning treatment, with one of the highest fecundities of 1,245,600 ± 552,117 eggs kg^-1^ bw obtained from females that were successfully induced.

Regarding egg quality of fertilised eggs incubated in 96 well plates, 96 ± 3% of fertilized eggs developed embryos and hatched indicating a high egg quality. Moreover, larvae survived as long as 13 days at 21°C without exogenous feeding which is considered a great improvement compared to the previous study by Ramos-Júdez *et al*. (2021), in which larvae survived no longer than 4 dph at 24 °C. Not only larvae survived longer without the application of external feeding, but also larvae reared using mesocosm conditions demonstrated the potential to develop until juveniles, indicating that eggs and larvae obtained after the induction of gametogenesis with rFsh and rLh could supply a hatchery. Indeed, larval development and growth in our study, using mesocosm rearing conditions, was very similar to other published studies either using intensive or extensive conditions (Abraham et al., 1999; Besbes et al., 2020; El-Gharabawy and Assem, 2006; Loi et al., 2020). Most of these studies, emphasized the effects of algal addition to the rearing tanks (green water technique) as the best way to optimize larval feeding on rotifers due to its effect not only in facilitating the contrast but also improving the nutritional state and / or health of the rotifers (Tamaru et al., 1994). Larval growth in size was similar to what Besbes *et al*. (2020) described, with larvae measuring 2.1 mm TL at hatching, and 2.96 mm TL at 4 dph, followed by a stagnated growth between 4 dph and 11 dph when the larvae reached 3.5 mm TL. Growth was accelerated from 13 dph to 35 dph when they reached 5 mm TL and then until day 39 when they reached almost 7 mm TL. The exponential growth recorded in the present study (Fig 14) was similar to that described by Besbes *et al*. (2020). Therefore, it was promising that the larvae reared in the mesocosms demonstrated similar grow and development as other studies on flathead grey mullet larvae indicating the potential for hatchery production of larvae from adults that had gametogenesis-induced with rGths.

Two critical periods have been described (Ignatius et al., 2017) during larval rearing of flathead grey mullet: one at days 2 - 3 post hatch due to the yolk sac resorption, a decrease in lipid reserves and an increase in specific gravity of the larvae, sinking to the bottom of the tank and gradually perish (Maslova, 2011), and the other, at 8-11 dph during swim bladder formation with an excessive inflation, especially in intensive (“artificial”) rearing systems (Nash et al., 1977; Nash and Kuo, 1975). These periods would justify the higher mortalities from 85 ± 18 % survival at 2 dph to 55 ± 17 % at 9 dph and 8 ± 11 % at 12 dph observed in larvae maintained in 96-well plates without exogenous feeding. Although survival was not the aim of the larvae rearing in mesocosms, the main mortality problem encountered was swimbladder hyperinflation. The larvae affected (see Fig 13G) remained floating in the water surface without moving and / or feeding on rotifers. This over inflation has been observed in other marine fish larvae such as in meagre (*Argyrosomus regius*) larvae (Vallés and Estévez, 2013), and is often associated with stressful conditions such as high light intensity, the use of long photoperiod, too early introduction of prey during larval rearing, or high larval density (Grotmol et al., 2005; Roo et al., 2010; Villamizar et al., 2011) that induces the larvae to gulp too much air in the water surface inducing the hypertrophy of the swim bladder.

## Conclusion

The approach described in the present study to induce oogenesis from previtellogenesis or early vitellogenesis to the completion of oocyte growth and spawning using single chain recombinant gonadotropins (rFsh and rLh) produced in CHO cells, offers replicability and guarantees a high success in spawning and high egg quality in flathead grey mullet. It was significant that in addition to inducing maturation from early gametogenesis through to the production of viable male and female gametes, the rGths have also induced the processes and cascade of hormones and pheromones that control reproductive behaviour and successful courtship in both females and males. From an applied point of view, the present protocol provides full control of reproduction with long-term weekly rGth administration. The protocol provided high fecundities from flathead grey mullet females (~ 1,700,000 eggs female^-1^), with fertilisation and hatching of ~ 50 % of the spawned eggs. These fecundities, indicate that the induction of 6 - 7 females (~1 kg) per season could permit a hatchery production of ~ 1 million fry, based on survivals reported in the literature. This would reduce the need of many breeders and the quantity of hormones used. Besides, the present protocol can probably be applied to develop out-of-season spawning and breeding programs.

## 6.0 Acknowledgements

Special thanks are due to Cristian Martínez Rodríguez, Alex Rullo Reverté, Esteban Hernández, Magda Monllaó and Sandra Molas for taking care of the fish and larvae and the technical help in samplings. Thanks also to Marta Sastre, Edgar Bertumeu and Noelia Gras for the help in steroids analysis. This study was funded by the Spanish Government, MINECO, project: RTI2018-094710-R-I00 coordinated by N.D. The participation of S.R. was funded, by a PhD grant from AGAUR (Government of Catalonia) co-financed by the European Social Fund.

